# Hydrophobic pocket engineering of arylmalonate decarboxylase expands its substrate scope towards the synthesis of the (*R*)-enantiomers of sterically hindered carboxylic acids

**DOI:** 10.64898/2026.05.07.723505

**Authors:** Elske van der Pol, Lisa-Marie Krammer, Johannes Eder, Dominik Gross, Roland C. Fischer, Kenji Miyamoto, Rolf Breinbauer, Robert Kourist

## Abstract

Arylmalonate decarboxylase (AMDase) stereoselectively converts disubstituted malonates to chiral carboxylic acids, but its substrate spectrum is very limited regarding the size of the smaller substituent. Inspired by the observation that (*S*)-selective AMDase variants also convert larger substrates, we unlocked the synthesis of the (*R*)-enantiomers of α-aryl and α-alkenyl *n-*butanoic and *n-*pentanoic acids, respectively, in exquisite enantiopurity.

---

With their high selectivity and mild reaction conditions, enzymes have established themselves as a widely used catalyst for the preparation of enantiomerically pure molecules.^1^ Limitations of the substrate scope of enzymes are a frequently encountered challenge, which can be addressed by enzyme engineering.^2,3^ The synthesis of α-substituted carboxylic acids is a typical example of the complementary strengths of chemical and enzyme catalysis. Direct alkylation of arylacetic acids using chiral auxiliaries is a practicable and widely used approach.^4,5^ Recently, direct stereoselective alkylation of arylacetic acids using chiral lithium amide as stereodirecting agent has been reported,^6^ giving access to a large number of α-alkyl carboxylic acids with enantiomeric excess ranging from 90-95%. A disadvantage is the requirement of the stoichiometric addition of an optically pure ligand. While various biocatalytic approaches for the discrimination of α-substituted carboxylic acids have been developed, including hydrolases and alcohol dehydrogenases,^7^ enantioselectivity is often unsatisfactory. Asymmetric decarboxylation of prochiral α,α-disubstituted malonic acids by bacterial arylmalonate decarboxylase (AMDase) gives access to α-aryl and α-alkenyl alkanoic acid derivatives such as non-steroidal anti-inflammatory drugs (NSAIDs),^8–12^ α-heterocyclic propionic acids^13,14^ in high yield and excellent stereoselectivity.^10,15^

The stereoselectivity of the enzyme could be switched by transplanting the catalytic Cys-residue to the opposite side of the substrate^16^ and the activity of the resulting (*S*)-selective AMDase variant was improved by focused directed evolution,^8,11,15,17^ which has been explained by the formation of a second hydrophobic pocket in the active site.^18^

AMDase accepts an impressive number of substrates having different larger substituents, with the only requirement that they should bear a delocalized π-electron system required for the stabilization of the nascent charge upon decarboxylation. In contrast, the acceptance of the second substituent is very restricted. AMDase accepts malonic acids with a hydrogen atom, a methyl, hydroxy, amine group, and halogen atoms.^19^ In their study on the first purification of the enzyme, Ohta and Miyamoto reported that 2-ethyl-2-phenyl malonate (**1a**) was inert to the enzyme.^20^ Recently, we could show that (*S*)*-*selective variants of AMDase having an extended hydrophobic pocket in the active site, such as AMDase V_43_I/G_74_C/A_125_P/V_156_L/M_159_L/C_188_G (‘AMDase ICPLLG’), accepted malonic acids with a larger second substituent and decarboxylated 2-ethyl-2-vinyl malonate (**2a**) to 2-ethylbut-3-enoic acid (**2b**).^21^ The excellent enantiomeric purity (>99 %ee) of the product demonstrated the outstanding capacity of AMDase to discriminate between the similarly sized ethyl and vinyl substituents.

In contrast, wildtype AMDase did not convert **2a** and showed only traces of **1b** after 20 h (**Fig. 1a**).^21^ On basis of this finding, we hypothesized that variants of the (*R*)-selective wildtype enzyme with an extended hydrophobic pocket could convert sterically hindered malonates with high stereoselectivity. Therefore, we prepared a series of α-aromatic and α-vinylic malonic acids with an ethyl and *n*-propyl group as second α-substituent (**1a-6a**) and investigated the activity and stereoselectivity of (*R*)-selective AMDase IPLL (V_43_I/A_125_P/V_156_L/M_159_L) with amino acid substitutions in the active site hydrophobic pocket (**Fig. 2A**). Our results confirm that the increased size and hydrophobicity of the active site lead to a wider substrate spectrum, thereby enabling access to optically pure α-aryl and α-alkenyl *n-*butanoic and *n-*pentanoic acids (**Fig. 1b**).

**Fig. 1.**
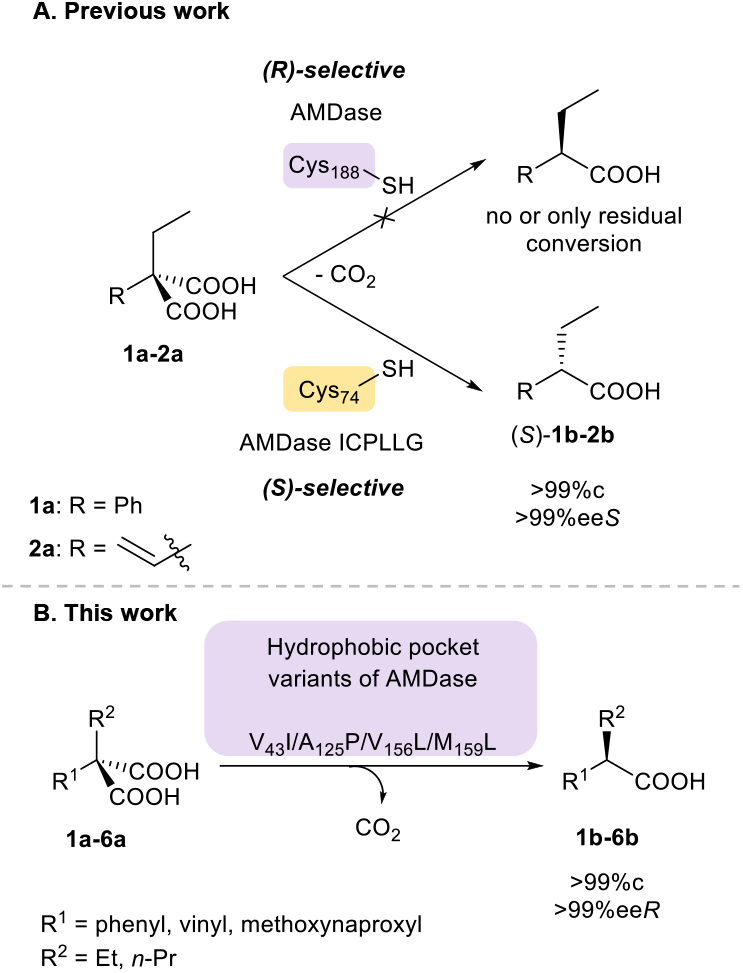
**A.**Previous work: Engineered (S)-selective AMDase accepts α-ethyl substrates, while (R)-selective AMDase WT does not. **B**. This work: (R)-selective AMDase variants with an engineered hydrophobic pocket convert α-ethyl and n-propyl substrates.

**Fig. 2.**
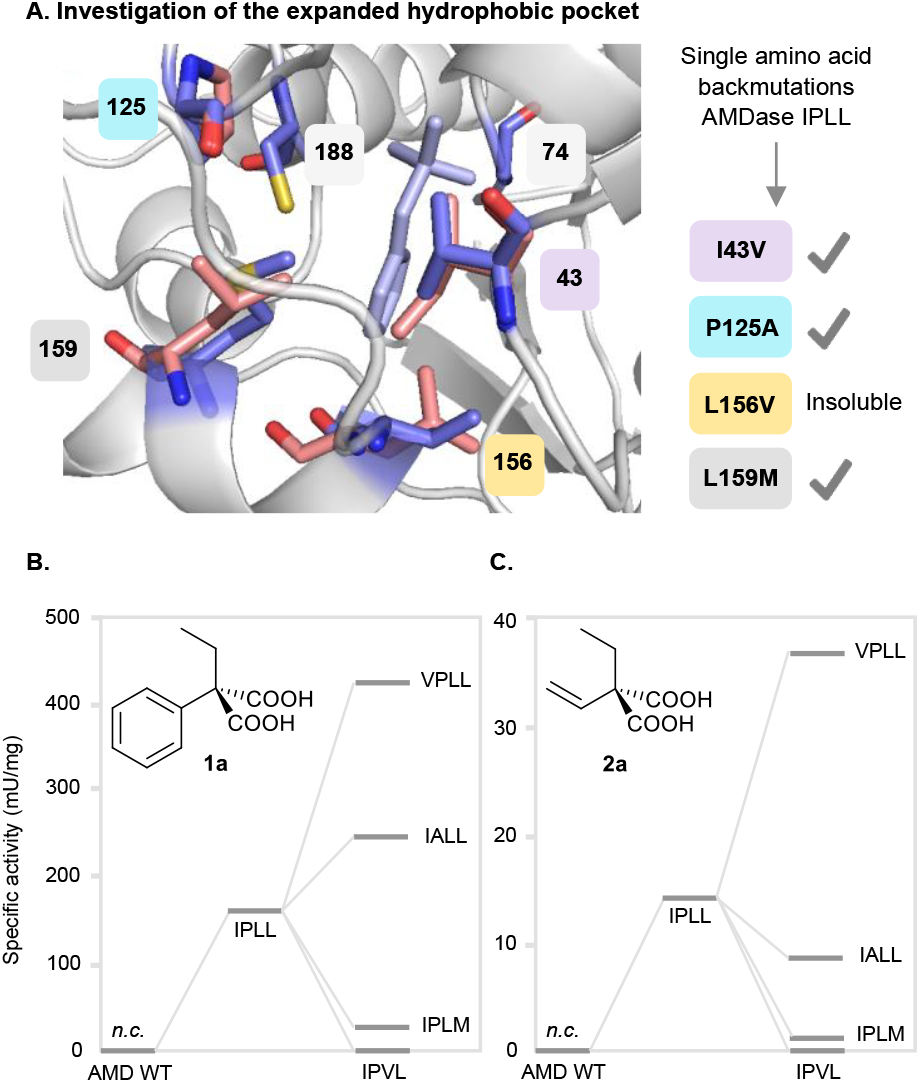
**A.**Active site of AMDase, with altered amino acids highlighted in purple (wildtype, PDB entry 3IP8) and light pink (IPLL variant). **B. C**. Specific activities of designed AMDase variants with substrates **1a-2a**. AMDase IPVL produced a low level of soluble enzyme and was barely active. See **Figure S3** for details, n=3.

In (*R*)-selective AMDase (C188), the mutations I_43_P_125_L_156_L_159_ increase activity towards aromatic α-methyl malonates.^11,15^ We were pleased to find that AMDase I_43_P_125_L_156_L_159_ also showed full conversion with **1a** and **2a** (**Fig 2B,C**), producing (*R*)-**1b** and (*R*)-**2b** in high optical purity (>98% ee). To test the effect of these single substitutions, I_43_P_125_L_156_L_159_ was back mutated by single-point mutations, resulting in four AMDase variants; AMDase V_43_P_125_L_156_L_159_, I_43_A_125_L_156_L_159_, I_43_P_125_V_156_L_159_, and I_43_P_125_L_156_M_159_ (**Fig. 2a**).

The amino acid substitution M_159_L leads to higher activity towards α-methyl malonic acids (**Fig S1**),^15,17^ therefore it is not surprising that I_43_P_125_L_156_L_159_ has also higher activity towards α-ethyl malonic acids than I_43_P_125_L_156_M_159_ (**Fig. 2B,C**). Unfortunately, soluble expression of AMD I_43_P_125_V_156_L_159_ was limited, and no specific activities could be measured for this variant. Reactions with cell-free extract demonstrated poor conversion. Variant I_43_A_125_L_156_L_159_ has higher activity towards **1a** than I_43_P_125_L_156_L_159_, whereas activity towards **2a** was insignificantly lower, showing that the contribution of this substitution has only a minor effect.

Surprisingly, back-mutation of I43 to valine (AMDase V_43_P_125_L_156_L_159_) further boosted the activity for both α-ethyl malonates (**Fig. 2B,C**), resulting in the fastest (*R*)-selective AMDase variant for the conversion of **2a** found so far (37.4 mU/mg, 2.7-fold improvement). In the liganded structure of AMDase G_74_C/C_188_S, the distance between the C_β_-atom of V43 and the C_α_-atom of the ligand phenyl acetate is only 5 Å. Substitution of isoleucine by the smaller valine residue, lacking one methyl group, likely reduces steric congestion in this region and thereby facilitates more productive substrate positioning, resulting in enhanced reaction rates towards α-ethyl-malonates. Although A-values for methyl and ethyl are similar, van der Waals volumes increase substantially, indicating that flexible alkyl substituents may impose steric constraints in confined enzyme active sites.^22^

The results of the backmutation show that the substitutions V_156_L and M_159_L contribute the strongest to the activity of I_43_P_125_L_156_L_159_. At position 43, L and V are possible, and at position 125, A and P. Not surprisingly, α-aromatic malonates react faster than α-vinylic (**Fig S1**), which has been observed before^15,21,23^ and can be attributed to the increased capacity to stabilize the nascent negative charge during the course of the reaction. For AMDase I_43_P_125_L_156_L_159_ and the faster AMDase V_43_P_125_L_156_L_159_, the measured optical purities are shown in **Table 1**. For (*R*)-**1b**, excellent ee’s were found for both variants, while for (*R*)-**2b**, reduced selectivity was observed. This is surprising as the (*S*)-selective AMDase ICPLLG well discriminates between the two substituents of **2a** (>99% ee). Back mutation of ICPLLG to AMDase VCPLLG resulted in a comparable positive trend in **2a** (**Fig. S1**). However, for **1a**, an adverse effect was observed. This can be rationalized by the altered binding mode of vinylic malonates compared to aryl malonates, in which the carboxylate is cleaved from the newly formed hydrophobic pocket.^19^ In this orientation, the residue at position 43 is part of the pocket accommodating the vinyl or ethyl substituent. Also, here, the substitution of isoleucine by the smaller valine residue reduces steric hindrance and allows for increasingly productive substrate positionings.

**Table 1.** Obtained enantiomeric excess (ee) of the chiral α-carboxylic acids produced by AMDase (on CFE). Aromatic acids 1b were derivatized to the corresponding methyl esters (Me-1b) using TMS-diazomethane. Experiments were performed in triplicate. n.d. = not determined. *first eluting peak.

| AMDase | (R)-Me-**1b** | (R)-**2b** | (+)-**3b** | (R)-**4b** | **5b** * |
| --- | --- | --- | --- | --- | --- |
| IPLL | >99% ± 0.0 | 94% ± 0.8 | 77% ± 0.0 | n.d. | n.d. |
| VPLL | >99% ± 0.0 | 96% ± 0.6 | 79% ± 0.7 | >99% ± 0.0 | >99% ± 0.0 |

We then proceeded to investigate other substrates posing specific challenges for the enzyme. For the well-accepted α-ethyl substituted substrate **2a**, we observed a 100-fold lower activity for both *(R)-* and *(S)-*selective AMDases compared to α-methyl-α-vinyl malonate (**Fig. S1**), which might be caused by steric reasons. To investigate the effect of the additional rotational degree of freedom, we examined **3a** with bridged α-groups, limiting the free rotation of the alkyl side group (**Fig. 3A,B**). Due to the conformationally restricted cyclohexenyl ring of **3a**, it should be considered that the 2-cyclohexene malonate adopts a half-chair conformation with one carboxylate in pseudoaxial and one in pseudoequatorial position.^24^ Typically, the axial substitutions are energetically less favored due to unfavorable interactions/steric clashes with other axial substituents and/or the p-orbitals of the unsaturated carbon-carbon bond. Therefore, bond cleavage of the α-carbon and the pseudoaxial carbonyl carbon would be favored. Subsequently, cleavage of the carboxylate introduces a partial negative charge on the α-carbon in the transition state. In contrast to **2a**, where the vinyl group has free rotation, for **3a** the partial negative charge can be stabilized through hyperconjugation of the aligned C=C bond. Both the conformational restrictions and the stabilizing effect would favor the acceptance of **3a** over **2a**.

**Fig. 3.**
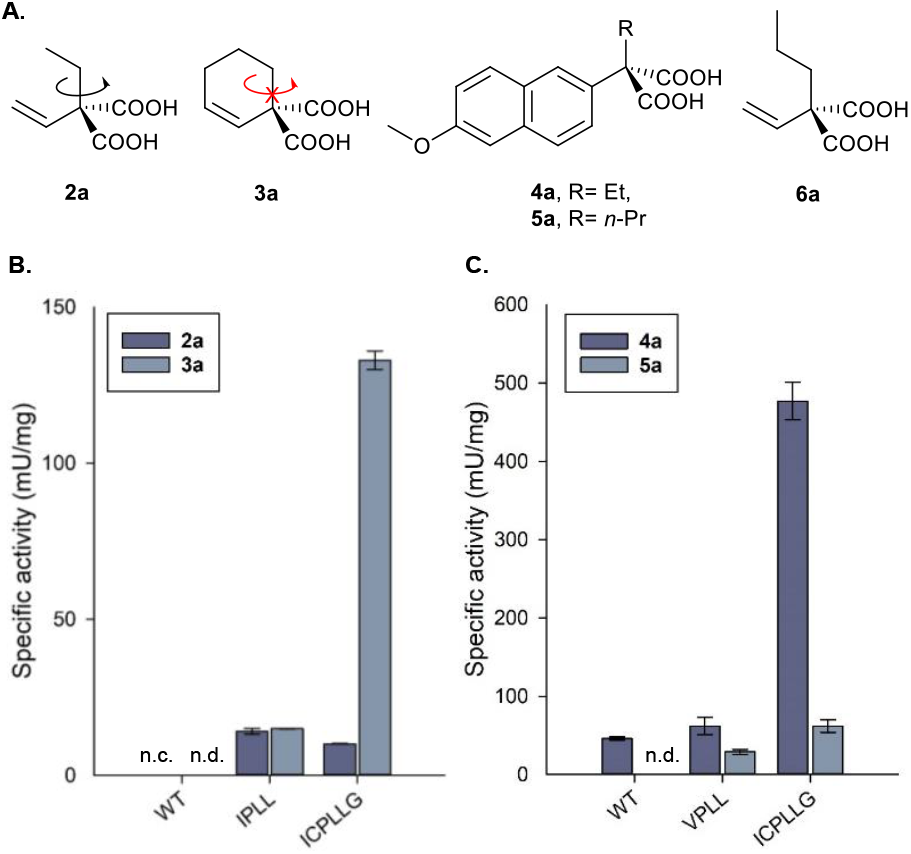
**A.**Various α-alkenyl and naproxen malonates. **B**. Specific activities of AMDase variants with **2a, 3a**. AMD WT does not convert **2a** (n.c.) **C**. Specific activities of AMDase variants with **4a** and **5a**. The conversion of **3a** and **5a** by AMD WT is too low to determine its specific activities (n.d.).

To investigate the impact of the G74C/C188G mutation on the specific activity, we determined the activity of the well-characterized (*R*)-selective AMDase IPLL and (*S*)-selective AMDase ICPLLG. For the AMDase IPLL, the rate for **3a** was similar to that for **2a** (approximately 15 mU/mg). In contrast, AMDase ICPLLG showed a 13-fold higher rate when comparing the strained malonate **3a** (133 mU/mg) to α-ethyl malonate **2a** (10 mU/mg) (**Fig. 3B**). Hence, we hypothesized that the limiting factor is not only the steric effect of the α-ethyl group, but also the optimal positioning of the reactive species, where the cleaved carboxylate, the α-carbon, and the cysteine would ideally be aligned at a 180-degree angle. While movement of the α-substituents is restricted for **3a**, the reaction coordinates in AMDase IPLL are not aligned at the optimal 180-degree angle required for the borderline concerted mechanism of AMDase variants towards alkenylic substrates,^21^ which would clarify the similar rates observed for both substrates by AMDase IPLL.

In contrast, when **2a** and **3a** adopt the same binding mode as 2-methyl-2-vinyl malonate in AMDase ICPLLG, the α-carbon would be perfectly aligned at a 180-degree angle with the leaving carboxylate and C74. This binding mode allows the reaction to be borderline concerted, with the protonation by C74 being partially rate-limiting. The free rotation of the α-substituents, being hampered and not obstructing protonation, further facilitates improved reactivity. Together, these effects might provide an explanation for the observed enhancement in the reaction rates obtained for **2a** and **3a** by AMDase ICPLLG (**Fig. 3B**). The stereoselectivity of the two (*R*)-selective AMDase variants towards **3a** was much lower than for **2a**. The small dialkyl malonates can bind in two inverse binding modes into the active site of AMDase, leading either to cleavage of the *pro*-*R* or the *pro*-*S* group, respectively.^21^ It can be easily imagined that due to the compact structure of **3a**, both binding modes have similar binding energies, making their discrimination very challenging.

We then investigated the synthesis of both enantiomers of optically pure ethyl-naproxen **4b**, which has recently been described as an effective aldo-keto reductase 1C3 inhibitor.^25^ Variants VPLL (*R*) and ICPLLG (*S*) were chosen as they had shown the highest activity towards α-ethyl substituted malonates (**Fig. 3C and 4**). AMDase ICPLLG exhibited the highest specific activity toward **4a**, while AMDase VPLL showed a ∼7-fold lower activity. Surprisingly, AMDase wildtype converted **4a**, albeit with 10-fold lower activity than AMDase VPLL. In comparison, the situation is reversed for the less sterically demanding naproxen malonate, which was converted 5-fold faster by the wildtype than by AMDase ICPLLG.^8^ This difference in the activity of wildtype and variants towards differently substituted α-aryl-α-ethyl substrates underlines that the effect of substitutions in the active site hydrophobic pocket of AMDase is hard to predict and highly substrate-specific.

**Fig. 4.**
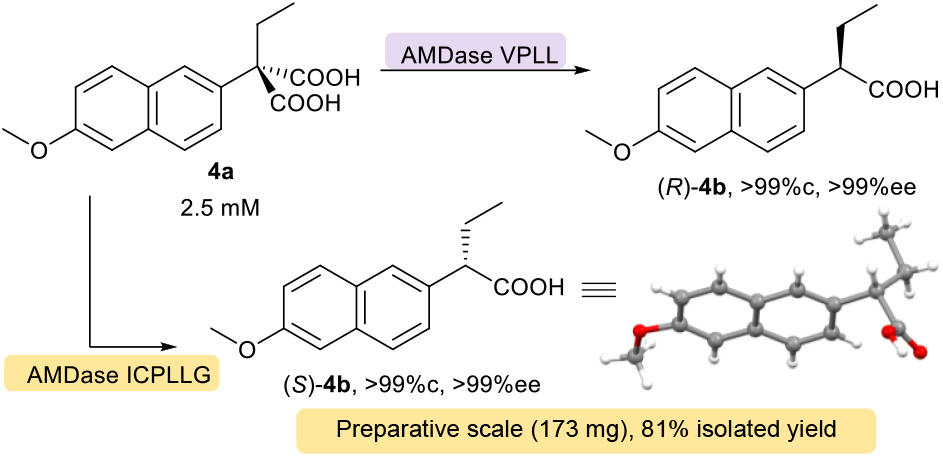
Conversion of **4a** by AMDase VPLL and ICPLLG. (S)-**4b** was produced on a preparative scale using (S)-selective AMDase ICPLLG.

Furthermore, we explored the acceptance of *n-*propyl-naproxen malonate (**5a**) and *n-*2-propyl-2-vinyl malonate (**6a**). While AMDase ICPLLG and VPLL indicated good acceptance of *n-*propyl-naproxen malonate (**Fig. 3C**), the vinyl compound demonstrated only limited conversion (39% and 19% for AMDase ICPLLG and VPLL, respectively, **Table S9**). We attribute this to the less effective stabilization of the evolving negative charge by the vinylic compound, combined with the steric hindrance of the *n*-propyl substituent. For the AMDase variants VPLL and ICPLLG, the specific activities toward **5a** were determined (**Fig. 3C**). Both variants exhibited markedly reduced rates for **5a** compared to those for **4a**, consistent with increased steric hindrance and greater rotational freedom of the *n*-propyl group. Overall, the engineered hydrophobic pocket of the variants ICPLLG and VPLL helps to overcome the limitations of the wildtype.

Both engineered AMDase variants were able to convert the naproxen derivatives **4a** and **5a** with excellent enantioselectivity (>99% ee) (**Table 1, Fig S36 and S39**), thereby providing access to both enantiomers of the aldo-keto reductase 1C3 inhibitor, α-ethyl naproxen. Recently, Adeniji and coworkers obtained the effective chiral acid **4b** by synthesizing the racemate, followed by chiral separation on preparative HPLC.^25^ Since AMDase ICPLLG and VPLL demonstrated the ability to produce the chiral acid with excellent purity, an enzymatic step in the synthesis process could be advantageous, potentially increasing the theoretical yield from 50% to 100%. With two stereocomplementary enzymes in hand, we wanted to demonstrate their scale-up potential. Using the more active (*S*)-producing enzyme, we synthesized (*S*)-**4b**. The preparative-scale biotransformation with AMDase ICPLLG using 173 mg of **4a** produced (*S*)-ethyl naproxen (**Fig. 4**) in 81% isolated yield, demonstrating the practicality to produce optically pure α-ethyl carboxylic acids with AMDase. We confirmed the absolute stereochemical configuration of **4b** obtained by AMDase ICPLLG-catalyzed decarboxylation *via* X-ray crystallography.

In conclusion, variation of active-site residues of AMDase, particularly the substitutions V_156_L and M_159_L, unlocked activity towards disubstituted malonic acids with an α-substituent larger than a methyl group. While wildtype decarboxylase has no or only very low activity towards these substrates, AMDase IPLL and variants thereof showed full conversion, enabling the enzymatic synthesis of α-aryl and α-alkenyl *n*-butanoic and *n*-pentanoic acids with very high stereoselectivity. It should be noted that point mutations in the hydrophobic pocket exert a strong and very substrate-specific effect on the conversion of these sterically hindered substrates, making it worth to investigate small sets of variants instead of a single AMDase variant for the conversion of sterically hindered malonic acids.

## Author contributions

E.v.d.P. conceptualized, coordinated, performed experiments, visualized, and prepared the manuscript. L.M.K. performed and analyzed the experiments with the naproxen malonates. She also prepared a part of the manuscript. J.E. generated the back-mutated AMDase variants and performed initial studies. D.G. performed the preparative synthesis of ethyl-naproxen malonate, the work-up of the preparative scales, and the crystallization. R.C.F. performed the X-ray diffraction and structure refinement. K.M. conceptualized and participated in discussions leading to the design of this study. R.B. conceptualized, provided funding, supervised, and prepared the manuscript. R.K. conceptualized, provided funding, supervised, visualized, and prepared the manuscript. All authors contributed to the data interpretation, reviewed, and approved the manuscript.

## Acknowledgements

We gratefully acknowledge financial support by the Austrian Science Fund (FWF) (Grant 10.55776/P34280) and NAWI Graz.

## Data availability

Experimental details, sequences, NMR spectra, (chiral) GC & HPLC chromatograms are available in the Supporting Information. Deposition Number 2545010 contains the supplementary crystallographic data for this paper. These data can be obtained free of charge via the joint Cambridge Crystallographic Data Centre (CCDC) and Fachinformationszentrum Karlsruhe Access Structures service.

