## Supplemental Information for "Hydrophobic pocket engineering of arylmalonate decarboxylase expands its substrate scope towards the synthesis of the (*R*)-enantiomers of sterically hindered carboxylic acids"

*Correspondence to:

#### 1. Additional Figures

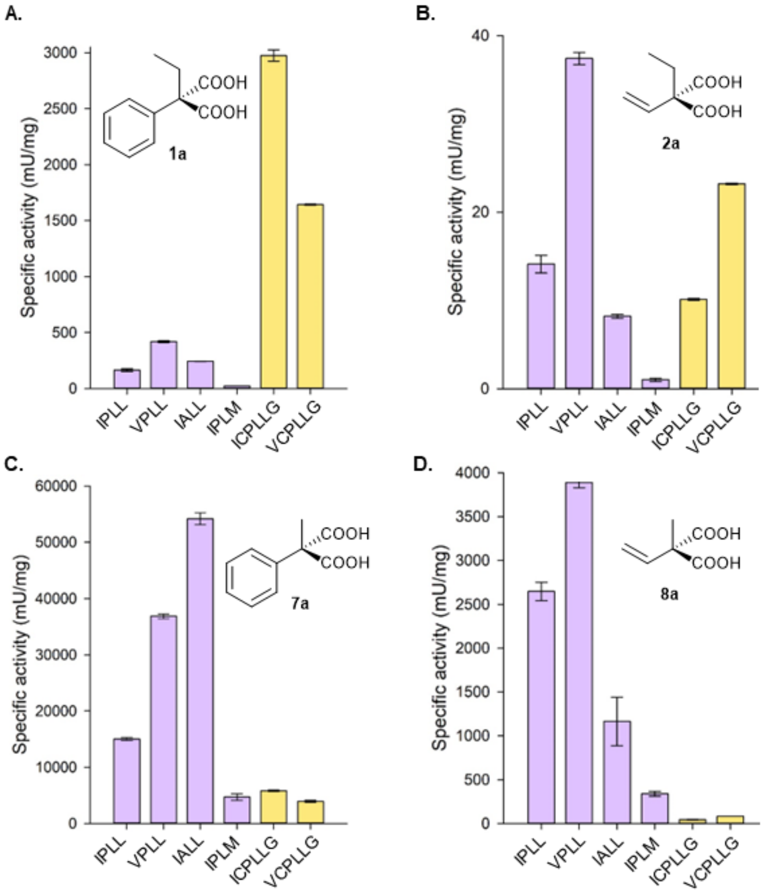

***Figure S1.*** *Specific activities of designed (R)- and (S)-selective AMDase variants with $\alpha$-methyl/$\alpha$-ethyl and vinyl/phenyl substrates (**1a**,**2a**, **7a**, and **8a**) on purified enzyme.*

***Table S1.*** *Obtained enantiomeric excess (ee) of the chiral $\alpha$-carboxylic acids produced by AMDase (on CFE). The ee's were determined by Chiral GC-FID after achieving full conversion (>99%) for **2a**, **3a**, and **8a**. Aromatic acids **1b** and **7b** were derivatized to the corresponding methyl esters using TMS-diazomethane. Experiments were performed in triplicate. n.d. = not determined. See sections 3.10 and 4 for further details.*

| AMD | Me-**1b** | **2b** | **3b** | Me-**7b** | **8b** |
|---|---|---|---|---|---|
| IPLL | >99% ± 0.0 (*R*) | 94% ± 0.8 (*R*) | 77% ± 0.0 (+) | >99% ± 0.1 (*R*) | 97% ± 1.6 (*R*) |
| VPLL | >99% ± 0.0 (*R*) | 96% ± 0.6 (*R*) | 79% ± 0.7 (+) | >99% ± 0.0 (*R*) | 98% ± 0.4 (*R*) |
| IALL | >99% ± 0.0 (*R*) | n.d. | n.d. | >99% ± 0.0 (*R*) | 98% ± 0.3 (*R*) |
| IPVL | 96% ± 0.0 (*R*) | n.d. | n.d. | >99% ± 0.0 (*R*) | 98% ± 0.3 (*R*) |
| IPLM | >99% ± 0.0 (*R*) | n.d. | n.d. | >99% ± 0.0 (*R*) | 99% ± 0.0 (*R*) |
| ICPLLG | >99% ± 0.0 (*S*) | >99% ± 0.0 (*S*) | 91% ± 3.0 (-) | 89% ± 1.3 (*S*) | 69% ± 0.3 (*S*) |
| VCPLLG | >99% ± 0.0 (*S*) | >99% ± 0.0 (*S*) | 87% ± 0.5 (-) | >99% ± 0.1 (*S*) | 58% ± 3.2 (*S*) |

#### 2. General Information

##### 2.1 Chemicals

All commercially available chemicals and solvents were purchased from Acros Organics, Alfa Aesar, Fisher, Fluka, Merck, Roth, Sigma-Aldrich, TCI, VWR, and used without further purification, unless otherwise stated.

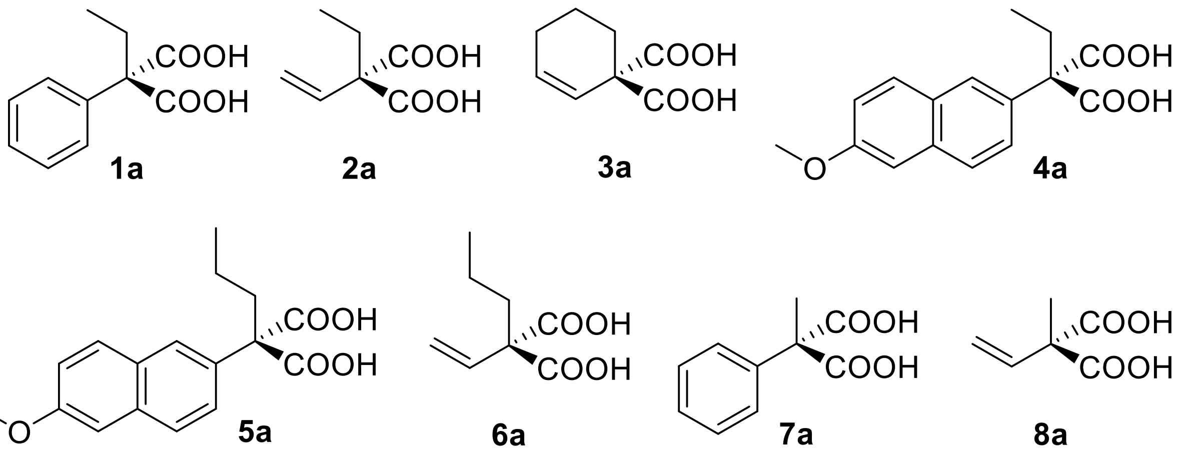

***Scheme S1.*** *Malonic acids used in this study.*

##### 2.2 Gas Chromatography

GC-FID: Chiral GC-FID analyses were performed on a Shimadzu Nexis GC-2030 device with a flame-ionization detector (FID) equipped with an Autosampler: AOC-20i/s 208. The separations were carried out on a Hydrodex-β-6TBDM column, (Chiral stationary phase, 25 m, 0.25 mm ID, 0.25 μm). The following isothermal methods were used: GC-FID_M1: Isothermal 103 °C for 35 min with a wash ramp 15 °C/min till 190 °C (for Me-**1b**), GC-FID_M2: Isothermal 100 °C for 30 min (for **2b** and **8b**), GC-FID_M3: Isothermal 159 °C for 10 min (for **3b**), and GC-FID_M4: Isothermal 100 °C for 35 min (for Me-**7b**). The temperature indicated is the column temperature.

##### 2.3 High Performance Liquid Chromatography

HPLC: Analytical HPLC measurements were performed on a Shimadzu HPLC SLC-40 equipped with a PDA detector (SDP-M40). All separations were carried out on a reversed-phase EC 250/4.6 NUCLEODUR C18 Pyramid (particle size: 5.0 μm, length: 250 mm, internal diameter: 4.6 mm) column. The following methods (***Table S2***) have been used to analyze the malonic acid substrates and their corresponding mono acid products:

***Table S2.*** *Parameters for the HPLC methods.*

| | **HPLC_M1 for 1a and 7a** | **HPLC_M2 for 2a and 3a** | **HPLC_M3 for 4a and 5a** | **HPLC_M4 for 6a** | **HPLC_M5 for 8a** |
|---|---|---|---|---|---|
| **Eluent** | 20 mM $H_3PO_4$ aq./$CH_3CN$ | 20 mM $H_3PO_4$ aq./$CH_3CN$ | 20 mM $H_3PO_4$ aq./$CH_3CN$ | 20 mM $H_3PO_4$ aq./$CH_3CN$ | 20 mM $H_3PO_4$ aq./$CH_3CN$ |
| **Ratio** | 70:30 | 80:20 | 60:40 | 70:30 | 85:15 |
| **Flow rate** | 1.0 mL/min | 1.0 mL/min | 1.0 mL/min | 1.0 mL/min | 1.0 mL/min |
| **Detector** | UV (254 nm) | UV (210 nm) | UV (230 nm) | UV (210 nm) | UV (210 nm) |
| **Column Temperature** | 40 °C | 40 °C | 40 °C | 40 °C | 40 °C |
| **Time** | 25 min | 25 min | 25 min (**4a**), 40 min (**5a**) | 20 min | 20 min |
| **Injection volume** | 5 μL | 5 μL | 2.5 μL | 2.5 μL | 5 μL |

#### 2.4 Chiral High Performance Liquid Chromatography

Method Chiral-HPLC: Analytical HPLC measurements were performed on a Shimadzu HPLC SLC-40 equipped with a PDA detector (SDP-M40). Chiral separations (for **4b** and **5b**) were carried out on a normal-phase ChiralPak IC column (particle size: 5.0 μm, length: 150 mm, internal diameter: 4.6 mm) with a mobile phase consisting of 0.05% TFA in *n*-heptane/ethanol (99:1%, *v*/*v*) at a flow rate of 0.5 mL/min, room temperature. Absorbance was monitored at 230 nm for 30 min, injection volume 1 μL.

#### 2.5 Nuclear Magnetic Resonance Spectroscopy

NMR spectra were recorded on a Bruker AVANCE III 300 spectrometer ($^1$H: 300.36 MHz; $^{13}$C: 75.53MHz) with an autosampler. Chemical shifts δ are referenced to the residual proton and carbon signal of the deuterated solvent ($CDCl_3$: δ = 7.26 ppm ($^1$H), 77.16 ppm ($^{13}$C); $CD_3OD$: δ

= 3.31 ppm ($^1$H), 49.00 ppm ($^{13}$C). Chemical shifts δ are given in ppm (parts per million) and coupling constants *J* in Hz (Hertz). Signal multiplicities are abbreviated as s (singlet), br s (broad singlet), d (doublet), dd (doublet of doublet), td (triplet of doublet), t (triplet), dt (doublet of triplet), q (quadruplet), p (pentet), and m (multiplet). Deuterated solvents for nuclear resonance spectroscopy were purchased from euriso-top®.

#### 2.6 Determination of Melting Points

Melting points were determined on a Mel-Temp® melting point apparatus from Electrothermal with an integrated microscopical support. They were measured in open capillary tubes with a mercury-in-glass thermometer and were not corrected.

#### 2.7 Determination of Optical Rotation

The specific optical rotation was determined on a Perkin-Elmer Polarimeter 341 with an integrated sodium vapor lamp. All samples were measured at the D-line of the sodium light (λ = 589 nm) in a 10 cm cell. Concentrations are given in g/100 mL. Each optical rotation measurement was performed ten times, and the mean value is reported.

#### 2.8 X-ray Diffraction

For single crystal X-ray diffractometry, all suitable crystals were covered with a layer of silicone oil. A single crystal was selected, mounted on a glass rod on a copper pin, and placed in the cold $N_2$ stream provided by an Oxford Cryosystems cryometer (T = 100 K), if not otherwise stated. XRD data collection was performed on a Bruker APEX II diffractometer with the use of Mo Kα radiation (λ = 0.71073 Å) from an IµS microsource and a CCD area detector. Empirical absorption corrections were applied using SADABS.[1] The structures were solved with the use of either direct methods or the Patterson option in SHELXS. Structure refinement was carried out using SHELXL.[2] CIF files were edited, validated, and formatted with the program OLEX2.[3] The space group assignments and structural solutions were evaluated using PLATON.[4] All non-hydrogen atoms were refined anisotropically. All hydrogen atoms were placed in calculated positions corresponding to standard bond lengths and angles using the riding model.

#### 2.9 Strains

The following strains have been used for cloning experiments and production of recombinant proteins, respectively; *E. coli* TOP10, genotype: F- mcrA Δ(mrr-hsdRMS-mcrBC) Φ80(lacZ)ΔM15 ΔlacX74 recA1 araD139 Δ(ara-leu)7697 galU galK rpsL(StrR) endA1 nupG, and *E. coli* BL21(DE3), genotype: F- ompT hsdSB ($r_B^- m_B^-$) gal dcm (DE3). Both strains have been purchased from Thermo Scientific.

#### 2.10 Plasmids

The following plasmids have been used: AMDase, AMDase IPLL, AMDase ICPLLG into the pET28(+) vector. All contain a kanamycin resistance marker, a *C-terminal* 6xHis-Tag, and an *NcoI/HindIII* cloning site. AMDase IPVL was ordered from *Twist Bioscience* into the pET-28a(+) vector with a C-terminus His-Tag. The codons were optimized for *E. coli*. AMDase VCPLLG has been reported in van der Pol et al.[5]

#### 2.11 DNA and amino acid sequence AMDase WT

<u>DNA sequence</u>

ATGGGCCAAATGCAACAGGCAAGCACCCCGACCATTGGTATGATCGTACCTCCCGCAGCTGGACTAGT
GCCAGCCGACGGCGCTCGTCTGTATCCGGATTTGCCGTTTATCGCGTCGGGTTTGGGCCTGGGATCCGT
GACTCCCGAAGGCTATGACGCGGTTATAGAAAGCGTGGTTGATCATGCTCGTCGCCTGCAAAAACAGG
GGGCAGCCGTTGTGTCTCTCATGGGGACCTCCTTGTCGTTCTACCGCGGCGCCGCTTTTAACGCGGCG
CTGACTGTCGCCATGCGTGAGGCTACTGGCCTGCCGTGTACTACCATGAGTACCGCCGTGCTAAACGGC
CTGCGTGCACTGGGCGTCCGGCGTGTGGCACTGGCGACCGCCTATATCGATGATGTTAATGAACGGCTT
GCAGCGTTTCTGGCGGAAGAATCCCTGGTACCGACAGGTTGTCGTAGCTTAGGCATTACCGGAGTAGA
AGCGATGGCTCGAGTGGATACCGCCACTCTGGTCGATCTGTGCGTCCGCGCCTTTGAAGCAGCACCAG
ATAGCGATGGGATTCTGTTGTCGTGTGGCGGTCTGCTTACCTTGGACGCAATCCCGGAAGTCGAGCGT
CGCCTGGGTGTGCCAGTCGTCTCAAGCAGTCCGGCGGGCTTTTGGGATGCGGTCCGTTTGGCTGGCG
GAGGCGCCAAAGCACGCCCGGGTTACGGCCGGCTTTTTGATGAGTCCGGTGGCAGCCACCATCACCA
TCACCATTAA

Amino acid sequence

MGQMQQASTPTIGMIVPPAAGLVPADGARLYPDLPFIASGLGLGSVTPEGYDAVIESVVDHARRLQKQGA
AVVSLMGTSLSFYRGAAFNAALTVAMREATGLPCTTMSTAVLNGLRALGVRRVALATAYIDDVNERLAAFL
AEESLVPTGCRSLGITGVEAMARVDTATLVDLCVRAFEAAPDSDGILLSCGGLLTLDAIPEVERRLGVPVVSS
SPAGFWDAVRLAGGGAKARPGYGRLFDESGGSHHHHHH

#### 2.12 Isolation of plasmid DNA

Cell material for plasmid isolation was either obtained from liquid culture (5-10 mL ONC) by centrifugation or from densely grown LB-agar plates. Commercially available GeneJET Plasmid Miniprep Kit (Thermo Scientific) or Wizard Plus SV Minipreps DNA Purification System (Promega) was used to isolate the plasmid DNA according to the manufacturer's instructions. Plasmid DNA was typically eluted in pure dd$H_2O$.

#### 2.13 Determination of DNA concentration

The concentration of isolated plasmid DNA or purified PCR products was determined spectrophotometrically with a NanoDrop 2000c spectrophotometer (Peqlab) at 260 nm. Additionally, the absorption at 280 and 230 nm was measured in parallel to assess the purity of the DNA sample.

#### 2.14 DNA sequencing and sequence analysis

Samples for Sanger sequencing were prepared according to the instructions provided by the company and sent either to GATC Biotech AG or Microsynth Austria GmbH. Respective sequencing primers were either available from the in-house primer collection or provided by the sequencing company. DNA sequences were aligned and analyzed with the online-tool Benchling.

#### 2.15 Buffer preparation

Reaction buffer (Tris HCl, 50 mM, pH 8): For the reaction buffer, 6.08 g Tris in 950 mL dd$H_2O$ and acidifying with 4M HCl to pH 8. The mixture was then filled up to 1000 mL with dd$H_2O$. For the purification, the reaction buffer was filtered under vacuum through a cellulose acetate filter (0.2 µm, Sartorius Stedim Biotech GmbH).

Binding buffer (Tris HCl (20 mM), NaCl (300 mM), Imidazole (20 mM), pH 7.4): For the binding buffer NaCl (17.52 g) and imidazole (1.36 g), and Tris (2.43 g) were dissolved in 930 mL dd$H_2O$. The solution was acidified to pH 7.4 with 4M HCl and filled up to 1000 mL. The buffer was filtered under vacuum through a cellulose acetate filter (0.2 µm, Sartorius Stedim Biotech GmbH).

Elution buffer (Tris HCl (20 mM), NaCl (300 mM), Imidazole (300 mM), pH 7.4): For the elution buffer NaCl (8.76 g) and imidazole (10.2 g), and Tris (1.21 g) were dissolved in 440 mL dd$H_2O$. The solution was acidified to pH 7.4 with 4M HCl and filled up to 500 mL. The buffer was filtered under vacuum through a cellulose acetate filter (0.2 µm, Sartorius Stedim Biotech GmbH).

Reverse-phase HPLC mobile phase ($H_3PO_4$, 20 mM): Concentrated Phosphoric acid (2.32 mL) was diluted in 2 L dd$H_2O$ and filtered under vacuum through a cellulose acetate filter (0.2 µm, Sartorius Stedim Biotech GmbH).

#### 2.16 BCA-assay

The total protein concentration of purified AMDase samples was determined using a PierceTM BCA Protein Assay Kit (Thermo Scientific). The samples were investigated as 1:10, 1:25, and 1:50 dilutions in triplicate each.

#### 2.17 Sodium Dodecyl - Polyacrylamide Gel Electrophoresis (SDS-PAGE)

For SDS-PAGE pre-cast gels (NuPAGE 4-12% Bis-Tris-Gel, Invitrogen) were used. The samples were prepared by diluting 4 µL CFE with 9 µL reaction buffer and adding 5 µL dye (NuPAGE LDS, Invitrogen) and 2 µL reducing agent (NuPAGE, Invitrogen). The samples were heated to 95 °C for 10 min before loading 17 µL each on the gel. As a standard, 4 µL PageRuler (ThermoScientific) protein ladder was used. The gels were run in Tris-MOPS buffer at 120 mA, max. 200 V for 50 min. Afterwards, the gels were stained with Coomassie blue overnight, destained and analyzed using the ImageJ software.

### 3. Biochemical procedures

#### 3.1 QuikChange mutagenesis

An adapted Q5 QuikChange protocol was used for single-point mutations of the AMDase IPLL variant. All required components were mixed together into a master mix with a total volume of 50 µL (***Table S3***). For negative control reactions, the polymerase was replaced with nuclease-free $H_2O$. The temperature program was selected based on the calculated annealing temperature of the primers, using no gradient and consisting of 26 cycles (***Table S4***).

***Table S3.*** *Master mix and temperature program for Q5 QuikChange mutagenesis PCRs.*

| **Component** | **Amount [µL]** | | **Step** | **Temperature** | **time** |
|---|---|---|---|---|---|
| 5x Q5 Reaction Buffer | 10 | | Initial denaturation | 98 °C | 30 s |
| dNTPs (10 mM) | 1 | | Denaturation | 98 °C | 10 s |
| Forward primer (10 µM) | 2.5 | | Annealing | $T_A$+1 °C | 20 s |
| Reverse primer (10 µM) | 2.5 | | Elongation | 72 °C | 4 min |
| Template DNA (undiluted) | 1 | | Extension | 72 °C | 5 min |
| 5x Q5 High GC enhancer | 10 | | Hold | 8 °C | ∞ |
| Nuclease free $H_2O$ | 22.5 | | | | |
| Q5 High Fidelity DNA Polymerase | 0.5 | | | | |
| Total Volume | 50 | | | | |

***Table S4.*** *List of primers used for Q5 QuikChange mutagenesis PCRs.*

| **Target mutation** | **Sequence** | $T_A$ | **GC-%** |
|---|---|---|---|
| I43V forward; ATC -> GTG | CCTGGGATCCgtgACTCCCGAAG | 67 °C | 65% |
| I43V reverse | CCCAAACCCGACGCGATA | 67 °C | 61 % |
| P125A forward; CCG -> GCC | ACTGGCGACCgccTATATCGATGATGTTAATGAACGGCTTGCAG | 72 °C | 50% |
| P125A reverse | GCCACACGCCGGACGCCC | 72 °C | 83% |
| L156V forward; CTG -> GTA | CATTACCGGAgtaGAAGCGCTGG | 64 °C | 57% |
| L156V reverse | CCTAAGCTACGACAACCTG | 64 °C | 53% |
| L159M forward; CTG -> ATG | ACTGGAAGCGatgGCTCGAGTGG | 70 °C | 61% |
| L159M reverse | CCGGTAATGCCTAAGCTACGAC | 70 °C | 55% |

##### 3.2 Transformation of chemo-competent *E. coli* TOP10 and *E. coli* BL21(DE3)

An aliquot of chemo-competent *E. coli* TOP10 or *E. coli* BL21(DE3) cells (50 µL) was thawed on ice, and DNA (5 µL, either isolated plasmid or QuikChange product) was added. The mixture was maintained on ice for 30 min, followed by a heat shock at 42 °C for 30 s. The mixture was cooled on ice for another 5 min. To promote growth, SOC media (950 µL) was added, and the cells were incubated at 37 °C and 350 rpm for 1 h. The cell suspension was concentrated by pelleting and partially discarding supernatant (850 µL). The leftover suspension was spread on LB-agar plates containing 40 µg/mL kanamycin and incubated overnight at 37 °C.

##### 3.3 Agarose gel electrophoresis

The size of DNA fragments was investigated using agarose gel electrophoresis. For gel preparation, 1% (w/w) agarose (LabQ Standard Agarose LE, LabConsulting) was dissolved in 1x TAE buffer by heating in a microwave until a homogeneous solution formed. After cooling to below 60 °C again, the DNA-intercalating dye (Atlas Clearsight DNA stain, BIOATLAS) was added. The mixture was poured into a small cast, a small-sized comb was inserted, and the gel was left to fully set. Samples (5 µL) were combined with 1 µL of 6x DNA Loading Dye (TriTrack, Thermo Scientific) and loaded onto the gel. A standard 1 kB GeneRuler DNA Ladder Mix (Thermo Scientific) served as the reference. The gels were run at 120 V and 400 mA for 35 minutes.

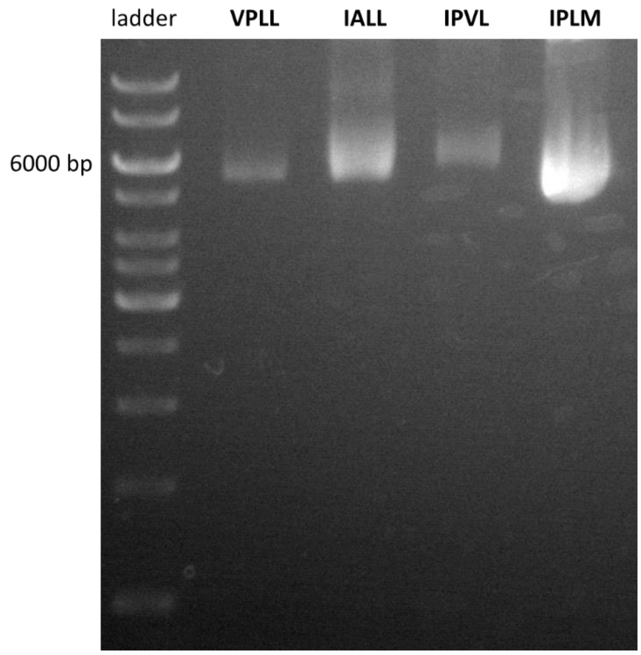

***Figure S2.*** *Agarose gel of the newly generated AMDase variants. A GeneRuler 1 kb DNA ladder was used for fragment size referencing.*

##### 3.4 Enzyme expression

AMD variants were expressed in *E. coli* BL21 (DE3). Overnight cultures were prepared from glycerol stocks in LB media (10 mL) with kanamycin (30 µg/mL) and were incubated at 37 °C, 120 rpm. The overnight cultures were used to inoculate using LB medium (250 mL) containing kanamycin (30 µg/mL). The cultures were incubated at 37 °C, 120 rpm until an $OD_{600}$ of 0.6-0.8 was reached. The expression was induced by adding IPTG (final concentration = 1 mM). The cultures were incubated at 28 °C, 120 rpm overnight. The cells were harvested by centrifugation at 4000 rpm, at 4 °C for 25 min. Subsequently, the pellets were resuspended in Tris HCl (50 mM, pH 8, 20 mL) and centrifuged at 4000 rpm at 4 °C for 25 min.

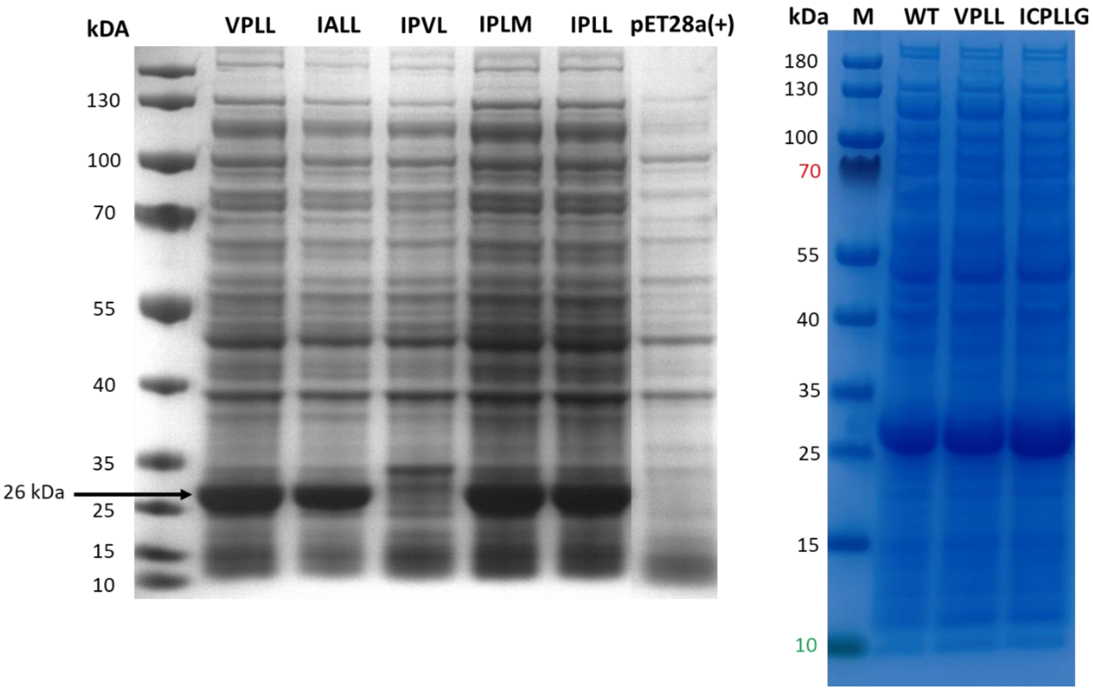

***Figure S3.*** *SDS-PAGE of E. coli BL21 DE3 cell-free extract after expression of new AMDase variants with AMDase IPLL as a reference. E. coli with an empty pET28a(+) vector served as a negative control. The expressed AMDase variants appear as thick bands with a size of 26 kDa. AMDase IPVL showed little soluble enzyme expression.*

##### 3.5 Enzyme purification

An AMDase pellet was defrosted on ice, whereafter the pellet was resuspended in Binding buffer (15 mL). The cells were sonicated on ice (Output control 50, Duty cycle 5, time: 2 min, followed by a 1 min break. Repeated 3x). The mixture was centrifuged at 13 000 rpm, at 4 °C for 25 min. The supernatant was filtered through a 0.45 µm non-pyrogenic sterile filter. Subsequently, the enzyme was purified using a Ni-Sepharose gravity column containing 2.5 mL "High Affinity Ni-Charged Resin" (GenScript). The CFE was loaded onto the column and the AMDase was allowed to bind by gently shaking on ice for 30 min. Afterwards, the flow through (FT) was collected and the column was washed with Binding buffer (5 mL, 2x). Subsequently, the enzyme was eluted with Elution buffer (2.5 mL, 2x).

##### 3.6 Buffer exchange

The Elution buffer was exchanged for the Reaction buffer (Tris HCl (50 mM, pH 8)) via a PD-10 Column (Sephadex™ G-25 M, Cytiva). The column was equilibrated with Reaction buffer (25 mL) before applying the Elution Fraction (EF) to the column. A maximum of 2.5 mL Elution

Fraction could be loaded onto the column simultaneously. The EF was entered into the packed bed completely, whereafter the Reaction buffer was used to elute the enzyme. Eluted fractions (1 mL) were collected into Eppendorf tubes. In total, six fractions were collected. Subsequently, the protein concentration was measured using Nanodrop (Protein A280) to detect the fractions with the highest enzyme concentrations. The fractions with the highest enzyme concentrations were combined, and a BCA assay was performed to measure the enzyme concentration more precisely.

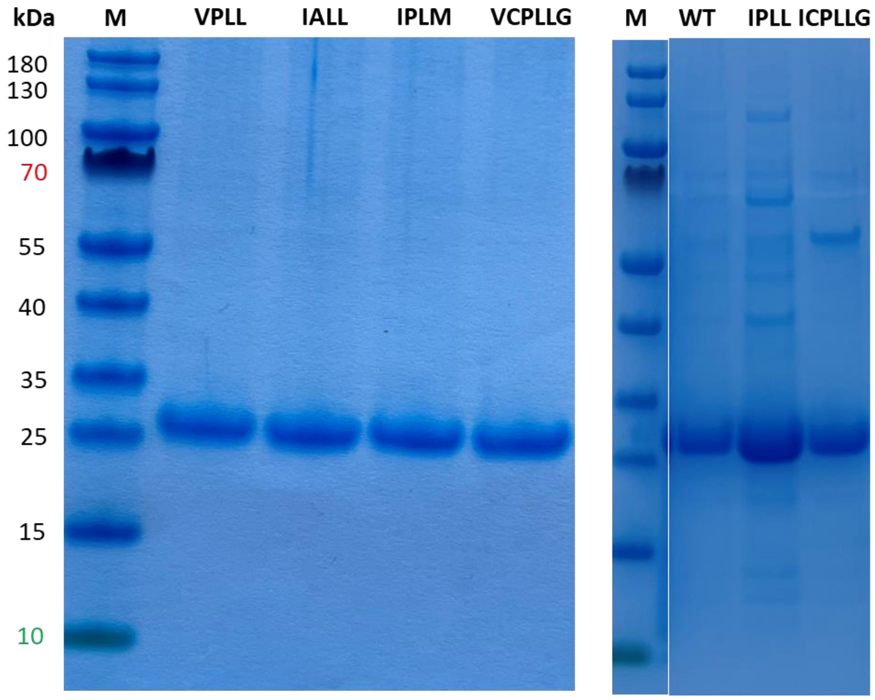

***Figure S4.*** *SDS-PAGE of purified AMDase variants used in this study.*

##### 3.7 AMDase Reaction rates for malonates 1a-3a, 7a, and 8a

Purified AMDase (700 μL) was added to substrate (700 μL) according to ***Table S5***, and the reaction was incubated at 30 °C, 600 rpm. Samples (150 μL) were taken and directly quenched with HCl (4M, 15 μL) at the following time points for each substrate:

**1a**: $t_0$, $t_5$, $t_{10}$, $t_{15}$, $t_{20}$, $t_{30}$, $t_{40}$, and $t_{50}$ minutes,

**2a**: $t_0$, $t_5$, $t_{10}$, $t_{15}$, $t_{20}$, $t_{30}$, $t_{40}$, and $t_{50}$ minutes,

**3a**: $t_0$, $t_5$, $t_{10}$, $t_{15}$, $t_{20}$, $t_{30}$, $t_{40}$, and $t_{50}$ minutes for AMD IPLL,

and $t_0$, $t_1$, $t_2$, $t_4$, $t_5$, $t_7$, and $t_{10}$, minutes for AMD ICPLLG.

**7a**: $t_0$, $t_1$, $t_2$, $t_4$, $t_5$, $t_7$, $t_{10}$, and $t_{15}$ minutes,

**8a**: $t_0$, $t_2$, $t_5$, $t_7$, $t_{10}$, $t_{15}$, $t_{20}$, and $t_{30}$ minutes.

***Table S5.*** *AMDase concentration (in mg/mL) used for the following substrate/variant combination. The substrate concentration used is indicated in mM.*

| | **1a**<br>**(10 mM)** | **2a**<br>**(2 mM)** | **3a**<br>**(2 mM)** | **7a**<br>**(10 mM)** | **8a**<br>**(1mM)** |
|---|---|---|---|---|---|
| **IPLL** | 0.5 | 0.75 | 0.5 | 0.005 | 0.05 |
| **VPLL** | 1.0 | 1.0 | n.d. | 0.005 | 0.05 |
| **IALL** | 1.0 | 1.0 | n.d. | 0.005 | 0.1 |
| **IPVL** | n.d. | n.d. | n.d. | n.d. | n.d. |
| **IPLM** | 1.0 | 1.0 | n.d. | 0.005 | 0.1 |
| **ICPLLG** | 0.01 | 1.0 | 0.4 | 0.005 | 0.1 |
| **VCPLLG** | 0.01 | 1.0 | n.d. | 0.005 | 0.1 |

***Table S6.*** *Rates AMDase variants (in mU/mg) on purified enzyme. Experiments were performed in triplicate (n = 3). The rates were analyzed by reverse-phase HPLC. Enzyme and substrate concentrations are described in* ***Table S5****.*

| **AMDase** | **1a** | **2a** | **3a** | **7a** | **8a** |
|---|---|---|---|---|---|
| **IPLL** | 164 ± 12 | 14.1 ± 1.0 | 15 ± 0.1 | 15047 ± 275 | 2647 ± 106 |
| **VPLL** | 418 ± 9 | 37.4 ± 0.7 | n.d. | 36867 ± 399 | 3891 ± 59 |
| **IALL** | 243 ± 2 | 8.2 ± 0.2 | n.d. | 54207 ± 1086 | 1164 ± 275 |
| **IPVL** | n.d. | n.d. | n.d. | n.d. | n.d. |
| **IPLM** | 23 ± 1 | 1 ± 0.2 | n.d. | 4720 ± 596 | 340 ± 28 |
| **ICPLLG** | 2973 ± 50 | 10 ± 0.1 | 133 ± 3 | 5820 ± 113 | 44 ± 3 |
| **VCPLLG** | 1643 ± 41 | 23 ± 0.1 | n.d. | 3960 ± 182 | 84 ± 1 |

#### 3.8 AMDase reaction rates for Naproxen malonates 4a-5a

Purified AMDase (700 µL) was added to substrate (700 µL) according to ***Table S7***, and the reaction was incubated at 30 °C, 600 rpm. Samples (100 µL) were taken, immediately quenched with 900 µL acetonitrile at the following time points for each substrate:

**4a**: $t_0$, $t_2$, $t_5$, $t_7$, $t_{10}$, $t_{15}$, $t_{20}$, and $t_{30}$ minutes,

**5a**: $t_0$, $t_2$, $t_5$, $t_7$, $t_{10}$, $t_{15}$, $t_{20}$, $t_{30}$, $t_{40}$, $t_{50}$, and $t_{90}$ minutes.

***Table S7.*** *AMDase concentration (in mg/mL) used for the following substrate/variant combination. The substrate concentration used is indicated in mM.*

| **AMDase** | **4a (2.5 mM)** | **5a (2.5 mM)** |
|---|---|---|
| **WT** | 0.75 | n.d. |
| **VPLL** | 0.5 | 0.75 |
| **ICPLLG** | 0.1 | 0.3 |

***Table S8.*** *Rates AMDase variants (in mU/mg) on purified enzyme. Experiments were performed in triplicate (n = 3). The rates were analyzed by reverse-phase HPLC. Enzyme and substrate concentrations are described in* ***Table S7****.*

| **AMDase** | **4a** | **5a** |
|---|---|---|
| **WT** | 46 ± 2 | n.d. |
| **VPLL** | 62 ± 11 | 29 ± 3 |
| **ICPLLG** | 477 ± 24 | 62 ± 8 |

##### 3.9 Conversion of malonates 5a-6a by AMDase

Cell pellets were resuspended in Tris-HCl buffer to a concentration of 75 mg wet cells/mL. The cells were sonicated on ice (Output control 50, Duty cycle 5, time: 2 min, followed by a 1 min break. Repeated 3x). The mixture was centrifuged at 13,000 rpm, at 4 °C for 25 minutes. The cell-free extract (CFE) was then mixed with substrate in a 1:1 ratio, resulting in a final substrate concentration of 2.5 mM. The reaction mixtures were incubated at 30 °C and 600 rpm, and samples were taken after 20 h. Samples containing *n*-propyl-naproxen (**5a**) were quenched by dilution 1:10 with acetonitrile, while those containing 2-*n*-propyl-2-vinyl malonate (**6a**) were quenched by adding 4 M HCl in a 10:1 ratio (sample: HCl).

***Table S9.*** *Conversion (%) of n-propyl malonates by AMDase variants on CFE (4.4 – 6.0 mg/mL) after 20 h. The conversions were measured via reverse-phase HPLC. Data represent mean values of three independent experiments (n=3). n.d. = not determined. n.c. = not converted.*

| **AMDase** | **5a (2.5 mM)** | **6a (2.5 mM)** |
| --- | --- | --- |
| **WT** | 43% ± 4 | n.c. |
| **VPLL** | n.d. | 19% ± 2 |
| **ICPLLG** | n.d. | 39% ± 4 |
| **Negative control (no enzyme)** | 0% ± 0 | 0% ± 0 |

##### 3.10 Determination of stereoselectivity (1b-3b, 7b, and 8b)

Cell pellets were resuspended in Tris-HCl buffer (50 mM, pH 8) to a concentration of 75 mg wet cells/mL. The cells were sonicated on ice (Output control 50, Duty cycle 5, time: 2 min, followed by a 1 min break. Repeated 3x). The mixture was centrifuged at 13,000 rpm, at 4 °C for 25 min. CFE containing AMDase (500 μL) was added to malonates **1a**-**3a**, **7a**, and **8a** (5 mM for **1a** and **7a**, 1 mM for **2a** and **3a**, and 2 mM for **8a**, 500 μL). The reactions were incubated at 30 °C, 600 rpm. After 3 h, another batch of CFE (500 μL) was added to reaction mixtures containing **2a**, **3a**, and **8a**. *Note: due to the inherent instability of malonates at elevated temperatures, complete conversion is required to obtain a reliable measurement of the enantiomeric excess. Incomplete conversion leads to spontaneous decarboxylation in the heated GC injector, forming a racemic reaction product and diluting the enantiomeric excess of the enzymatic product.*

At t = 20 h, the reactions were analyzed using reverse-phase HPLC. Reactions that fully converted the malonates into the corresponding chiral carboxylic acids were selected for chiral GC measurements. To the incomplete reactions, AMDase IALL, IPVL, and IPLM with **2a** and **3a**, freshly lysed CFE (500 μL) was added. The reactions were incubated at 30 °C, 600 rpm, 15 h. Afterward, the reactions were analyzed again using reverse-phase HPLC. Despite efforts to achieve complete conversion of the malonate, traces of malonate were still observed. These reactions were excluded from the determination of enantiomeric excess.

To prepare the samples for chiral GC measurements, the reaction samples (500 µL) were first quenched by the addition of HCl (50 µL, 4M). Subsequently, the compounds were extracted using EtOAc (300 µL) and centrifuged for 10 min at 13,000 rpm and 4 °C. The organic layer was dried over anhydrous $Mg_2SO_4$ and centrifuged (1 min, 13,000 rpm, 4 °C). Chiral carboxylic acids **2a**, **3a**, and **8a** were measured directly using chiral GC, while **1b** and **7b** were derivatized beforehand.

Derivatization of **1b** and **7b**

To the dried organic phase (300 µL), MeOH (100 µL) and $TMSCHN_2$ (25 µL) were added. After 20 min, the reaction was quenched by the addition of AcOH (2.5 µL). The sample was dried using an argon flow until completely dry. Subsequently, EtOAc (200 µL) was added, and the sample was measured by chiral GC. *Note: complete conversions are not required for malonates **1a** and **7a**, as their corresponding dimethyl malonates are stable under elevated temperatures.*

##### 3.11 Determination of stereoselectivity (4b-5b)

Purified AMDase (0.7 mg/mL) was added to **4a** or **5a** (5 mM, final substrate concentration was 2.5 mM). The reactions were incubated at 30 °C, 600 rpm, 20 h. To prepare the samples for chiral HPLC measurements, the reaction samples (300 µL) were first quenched by the addition of HCl (20 µL, 4M). Subsequently, the compounds were extracted using EtOAc (300 µL) and centrifuged for 10 min at 13,200 rpm and 4 °C. The organic layer was dried over anhydrous $Mg_2SO_4$ and centrifuged (1 min, 13200 rpm, 4 °C) and used for chiral HPLC measurements.

##### 3.12 Sample preparation for reverse-phase HPLC

After centrifugation for 1 h, 13,000 rpm at 4 °C, the samples were measured via HLPC (HPLC_M1 for compounds **1a** and **7a**, HPLC_M2 for compounds **2a** and **3a**, HPLC_M3 for compounds **4a** and **5a**, HPLC_M4 for compounds **6a**, and HPLC_M5 for compound **8a**). Because of the stronger UV absorption of the malonates, substrate depletion for **2a**, **3a**, and **8a** was plotted as a function of time. Only linear points were taken, and the slope was calculated.

#### 3.13 Preparative scale decarboxylation with AMDase ICPLLG of 2-cyclohexene-1,1-dicarboxylic acid (3a)

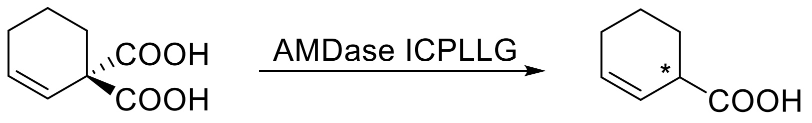

**3a**
4 mM
100 mg

(-)-**3b**
51 mg
69% isolated yield

<u>Expression of AMD ICPLLG</u>

AMD ICPLLG was expressed in *E. coli* BL21. Overnight cultures were prepared from glycerol stocks in LB media (100 mL) with kanamycin (30 µg/mL) and were incubated at 37 °C, 120 rpm. The overnight cultures were used to inoculate 800 mL of LB medium containing kanamycin (30 µg/mL). The cultures were incubated at 37 °C, 120 rpm until $OD_{600}$ was 0.6-0.8. The expression was induced by adding IPTG (final concentration = 1 mM). The cultures were incubated at 28 °C, 120 rpm overnight. The cells were harvested by centrifugation at 4000 rpm, at 4 °C for 25 min. Subsequently, the pellets were washed with Tris HCl (50 mM, pH 8, 20 mL), followed by centrifugation at 4000 rpm, at 4 °C for 30 min.

<u>Biotransformation Preparative scale</u>

The cells were resuspended in Tris HCl (50 mM, pH 8, 75 mg/mL of wet cell mass) and sonicated on ice as mentioned before for 2 min, followed by a 1 min break (3x). The CFE was centrifuged at 12,000 rpm, 4 °C for 25 min. AMDase ICPLLG CFE (19 mL) was combined with 2-cyclohexene-1,1-dicarboxylic acid (**3a**) (19 mL, 8 mM) in a 50 mL Falcon tube (4x) and the reactions were incubated at 30 °C, 600 rpm. After 16 h, the reaction was monitored via HPLC, which showed complete conversion of **3a**.

The reaction was acidified with HCl (4 M, 4 mL per Falcon tube), followed by centrifugation at 9,000 rpm, 30 min. EtOAc (150 mL) was added to the combined supernatant, and the mixture was centrifuged at 5,000 rpm for 7 min. The organic phase was washed with brine (50 mL), dried over $Na_2SO_4$, and filtered. The crude was concentrated under reduced pressure.

The crude product was purified via flash column chromatography (10 g $SiO_2$, 15.0 cm x 1.0 cm, *n*-pentane:$Et_2O$ = 6:1 to 3:1 (v/v) + 0.25vol% $HCO_2H$) affording the desired product as a colorless liquid (51 mg, 69%).

$C_7H_{10}O_2$ [126.16 g·mol$^{-1}$]

$R_f$ = 0.40 (cyclohexane:EtOAc = 4:1 (v/v) + 0.25vol% $HCO_2H$, $KMnO_4$)

$[\alpha]_D^{20}$ = -108.4 (c = 2.52, $CH_2Cl_2$)

$^1H$ NMR (300 MHz, $CDCl_3$) δ 10.30 (bs, 1H), 5.96 – 5.83 (m, 1H), 5.77 (m, 1H), 3.28 – 3.00 (m, 1H), 2.03 (m, 2H), 1.98 – 1.83 (m, 2H), 1.82 – 1.51 (m, 2H) ppm.

$^{13}C$ NMR (76 MHz, $CDCl_3$) δ 181.2, 130.3, 123.8, 41.0, 25.2, 24.7, 20.8 ppm.

#### 3.14 Preparative scale decarboxylation with AMDase ICPLLG of ethyl-naproxen malonate (4a)

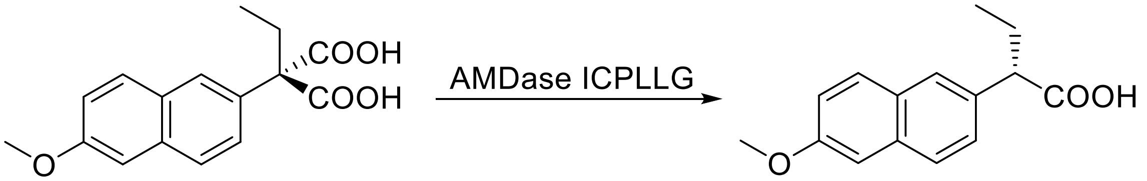

**4a**
2.5 mM
173 mg

(*S*)-(+)-**4b***
118 mg
81% isolated yield

**The stereochemistry of (S)-**4b** was determined via a crystal structure, which is described in the next chapter.*

AMDase ICPLLG was expressed, and CFE was prepared as described in the previous subchapter. AMDase ICPLLG CFE (15 mL) was combined with ethyl-naproxen malonate (**4a**) (15 mL, 5 mM) in a 50 mL Falcon tube (8x). The reaction mixture was then incubated at 30 °C at 600 rpm. Upon completion of the reaction, as confirmed by HPLC analysis after 2 h, the mixture was stored at -20 °C. Subsequently, the defrosted turbid mixture was centrifuged at 12,000 rpm for 30 min, 4 °C, and the supernatant was separated from the pellet.

The reaction mixture was extracted with EtOAc (3x 100 mL). The combined organic layers were dried over $Na_2SO_4$, filtered, and concentrated under reduced pressure. The crude product was purified via flash column chromatography (12 g $SiO_2$, 14.5 cm x 1.2 cm, cyclohexane:EtOAc = 2:1 to 3:2 (v/v)) affording the desired product as an off-white solid (118 mg, 84%).

$C_{15}H_{16}O_3$ [244.29 g·mol$^{-1}$]

$R_f$ = 0.27 (cyclohexane:EtOAc = 2:1 (v/v), UV, CAM)

$[\alpha]_D^{20}$ = +61.8 (c = 1.06, EtOAc)

$^{1}H$ NMR (300 MHz, $CDCl_3$) δ 10.27 (s, 1H), 7.71 (d, $^{3}J_{HH}$ = 8.4 Hz, 3H), 7.43 (d, $^{3}J_{HH}$ = 8.5 Hz, 1H), 7.15 (m, 2H), 3.91 (s, 3H), 3.61 (t, $^{3}J_{HH}$ = 7.6 Hz, 1H), 2.33 – 1.78 (m, 1H), 0.95 (t, $^{3}J_{HH}$ = 7.3 Hz, 1H) ppm.

$^{13}C$ NMR (76 MHz, $CDCl_3$) δ 180.6, 157.8, 134.0, 133.6, 129.4, 129.0, 127.3, 127.0, 126.6, 119.1, 105.7, 55.4, 53.4, 26.3, 12.2 ppm.

#### 4. Crystal Data and Structure Refinement

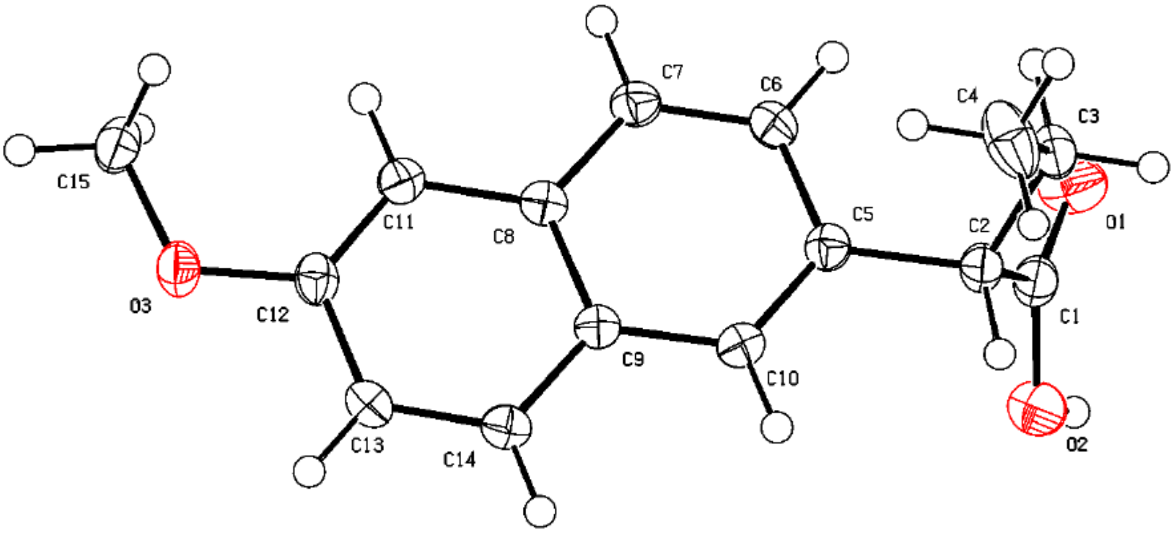

***Figure S5.*** *Molecular structure of (+)-ethyl naproxen (**4b**) obtained by a preparative scale reaction with AMDase ICPLLG. AMDase ICPLLG yields (S)-ethyl naproxen, (S)-**4b**.*

| | |
|---|---|
| Compound | (+)-**4b** |
| Empirical formula | $C_{15}H_{16}O_3$ |
| Formula weight [g·mol$^{-1}$] | 244.28 |
| Temperature [K] | 100.0(2) |
| Crystal system | monoclinic |
| Space group | $P2_1$ |
| a [Å] | 7.87968(9) |
| b [Å] | 5.68221(7) |
| c [Å] | 14.07726(19) |
| α [°] | 90 |
| β [°] | 91.1010(11) |
| γ [°] | 90 |
| Volume [Å$^3$] | 630.179(13) |
| Z | 2 |
| $\rho_{calc}$ [g·cm$^{-3}$] | 1.287 |
| μ [mm$^{-1}$] | 0.721 |
| F000 | 260.0 |
| Crystal size [mm$^3$] | 0.13 × 0.09 × 0.06 |
| Radiation | Cu Kα (λ = 1.54184) |
| 2Θ range for data collection [°] | 6.28 to 159.512 |
| Index ranges | -10 ≤ h ≤ 10, -7 ≤ k ≤ 7, -16 ≤ l ≤ 17 |
| Reflections collected | 32677 |
| Independent reflections | 2704 [$R_{int}$ = 0.0547, $R_{sigma}$ = 0.0199] |
| Data/restraints/parameters | 2704/1/167 |
| Goodness-of-fit on $F^2$ | 1.073 |
| Final R indexes [I ≥ 2σ(I)] | $R_1$ = 0.0317, $wR_2$ = 0.0836 |
| Final R indexes [all data] | $R_1$ = 0.0323, $wR_2$ = 0.0839 |
| Largest diff. peak/hole / e [Å$^{-3}$] | 0.21/-0.17 |
| Flack parameter | -0.06(10) |

### 5. Chiral GC chromatograms

The conversions of **1a**, **2a**, **3a**, and **8a** by AMDase ICPLLG, **2a** and **8a** by AMDase VCPLLG, and **8a** by AMDase IPLL have been reported by van der Pol et al.[5]

#### 5.1 Methyl-2-phenyl butanoate (Me-1b)

<u>Racemic methyl 2-phenyl butanoate (Me-**1b**)</u>

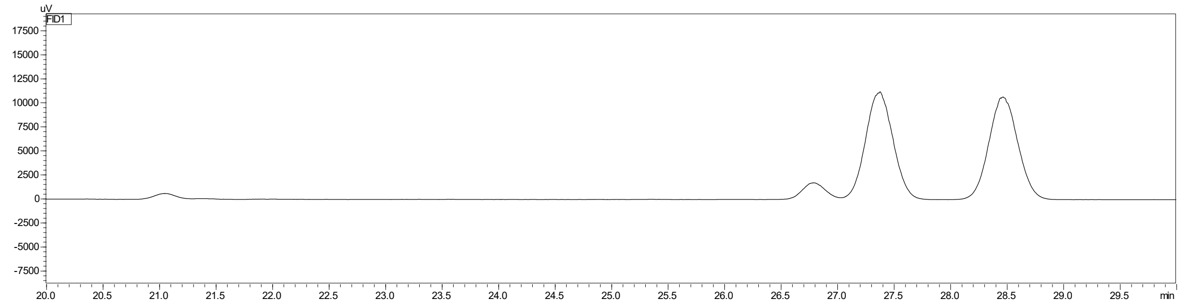

***Figure S6.*** *Chiral GC chromatogram of commercially available racemic methyl 2-phenyl butanoic acid after derivatization with TMSCHN$_2$ (Me-**1b**). Retention times of (S)-Me-**1b** = 27.3 min and (R)-Me-**1b** = 28.5 min. Method: GC-FID_M1. Retention times differ due to column exchange.*

<u>Conversion of **1a** by AMDase IPLL</u>

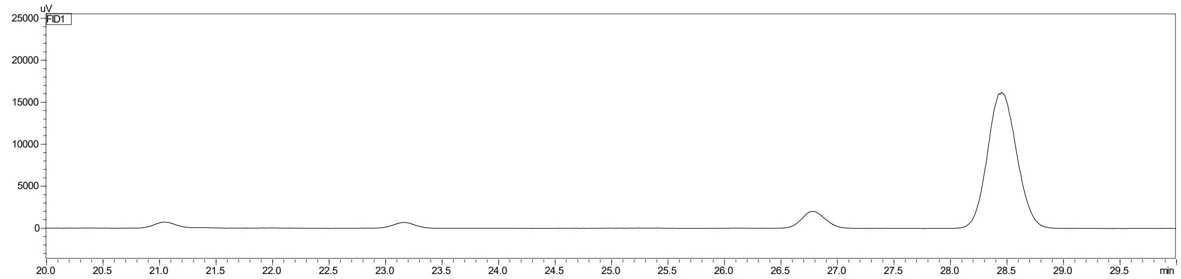

| **Peak** | **Retention time (min)** | **Area** |
|---|---|---|
| (*S*)-Me-**1b** | 27.2 | *not detected* |
| (*R*)-Me-**1b** | 28.5 | 288770 |

***Figure S7.*** *Chiral GC chromatogram of methyl 2-phenyl butanoate (Me-**1b**). **1b** was obtained from the AMDase IPLL-catalyzed conversion of 2-ethyl-2-phenyl malonate (**1a**). Method: GC-FID_M1. >99% ± 0.0 ee (R). Retention times differ due to column exchange.*

Racemic methyl 2-phenyl butanoate (Me-**1b**)

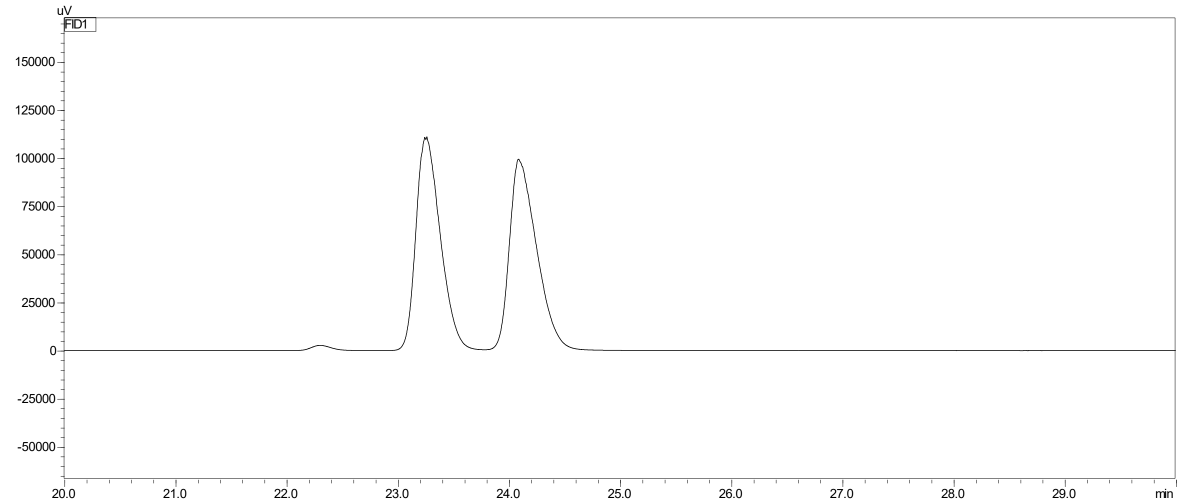

***Figure S8.*** *Chiral GC chromatogram of commercially available racemic methyl 2-phenyl butanoic acid after derivatization with TMSCHN*$_2$ *(Me-**1b**). Retention times of (S)-Me-**1b** = 23.3 min and (R)-Me-**1b** = 24.2 min. Method: GC-FID_M1.*

Conversion of **1a** by AMDase VPLL

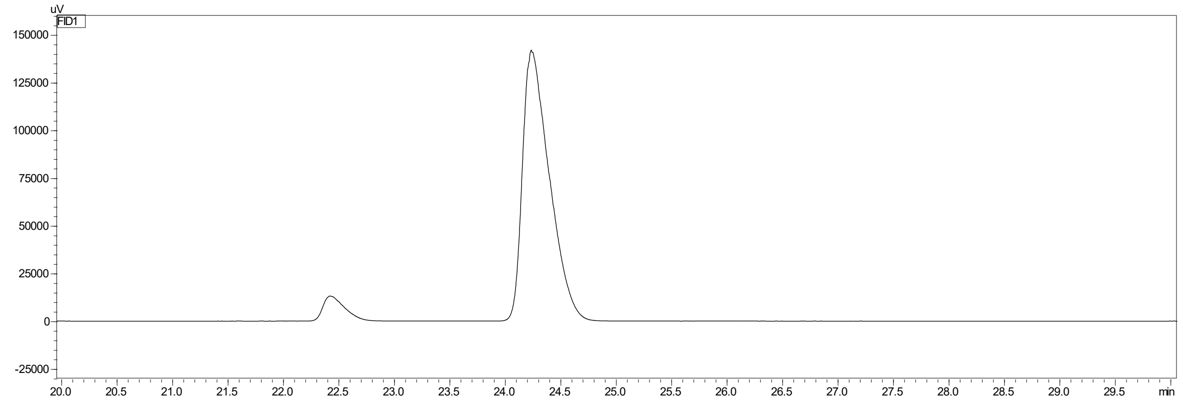

| **Peak** | **Retention time (min)** | **Area** |
|---|---|---|
| (*S*)-Me-**1b** | 23.4 | *not detected* |
| (*R*)-Me-**1b** | 24.2 | 2390663 |

***Figure S9.*** *Chiral GC chromatogram of methyl 2-phenyl butanoate (Me-**1b**). **1b** was obtained from the AMDase VPLL-catalyzed conversion of 2-ethyl-2-phenyl malonate (**1a**). Method: GC-FID_M1. >99% ± 0.0 ee (R).*

Conversion of **1a** by AMDase IALL

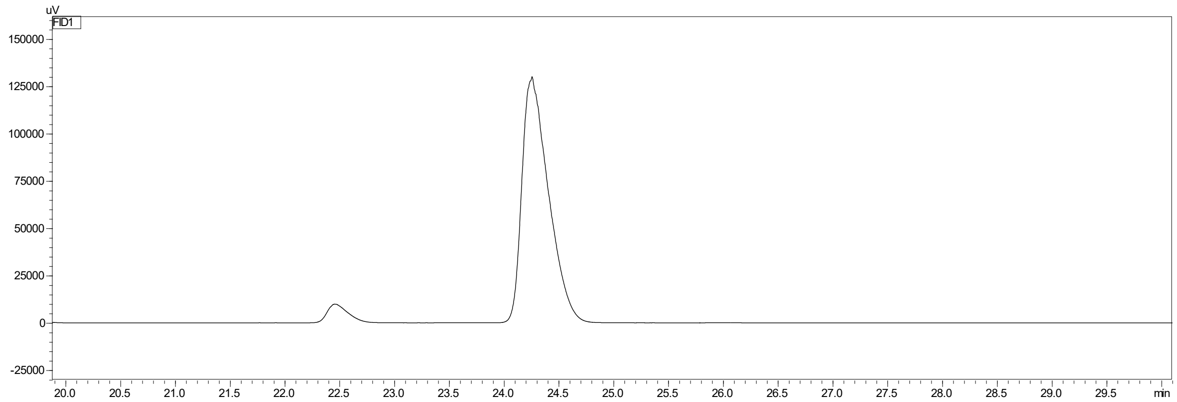

| Peak | Retention time (min) | Area |
| --- | --- | --- |
| (*S*)-Me-**1b** | 23.4 | *not detected* |
| (*R*)-Me-**1b** | 24.2 | 2180143 |

***Figure S10.*** *Chiral GC chromatogram of methyl 2-phenyl butanoate (Me-**1b**). **1b** was obtained from the AMDase IALL-catalyzed conversion of 2-ethyl-2-phenyl malonate (**1a**). Method: GC-FID_M1. >99% ± 0.0 ee (R).*

Conversion of **1a** by AMDase IPVL

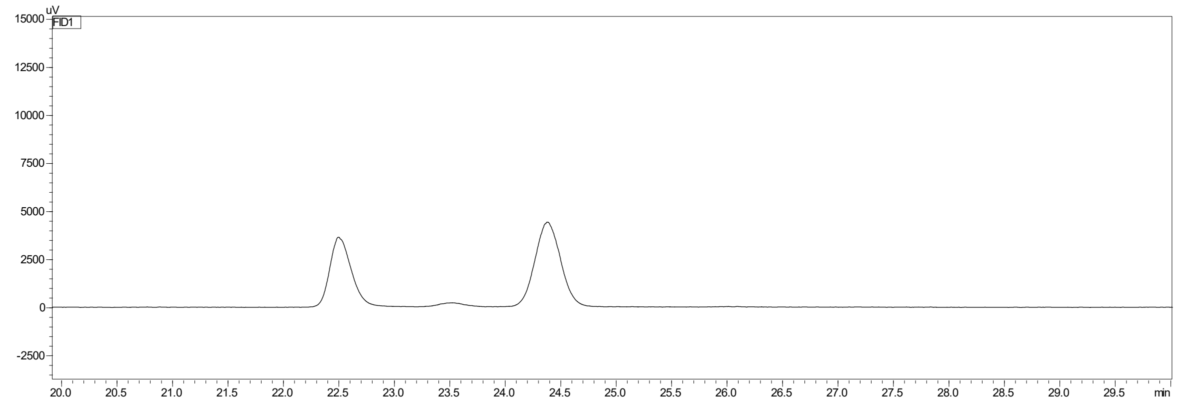

| Peak | Retention time (min) | Area |
| --- | --- | --- |
| (*S*)-Me-**1b** | 23.4 | 1495 |
| (*R*)-Me-**1b** | 24.2 | 69949 |

***Figure S11.*** *Chiral GC chromatogram of methyl 2-phenyl butanoate (Me-**1b**). **1b** was obtained from the AMDase IPVL-catalyzed conversion of 2-ethyl-2-phenyl malonate (**1a**). Method: GC-FID_M1. 95.8% ± 0.0 ee (R).*

Conversion of **1a** by AMDase IPLM

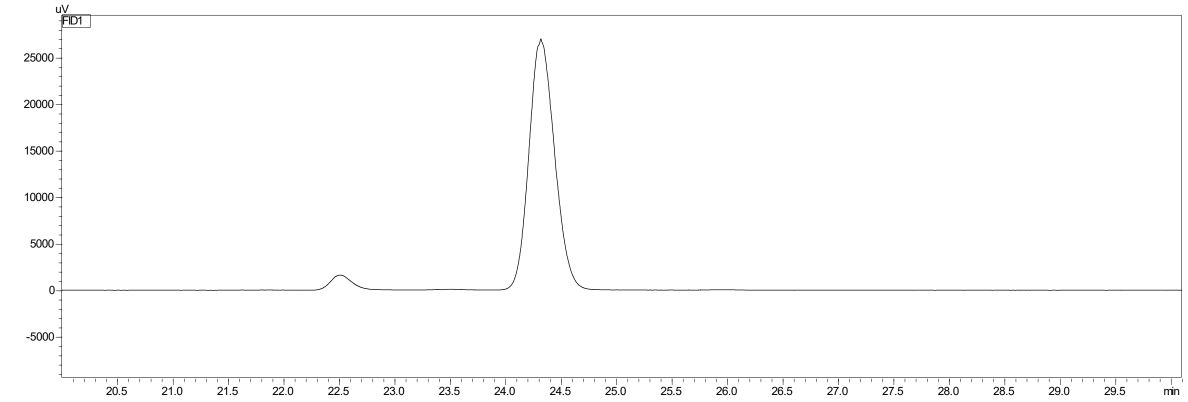

| Peak | Retention time (min) | Area |
|---|---|---|
| (*S*)-Me-**1b** | 23.4 | *not detected* |
| (*R*)-Me-**1b** | 24.2 | 430982 |

***Figure S12.*** *Chiral GC chromatogram of methyl 2-phenyl butanoate (Me-**1b**). **1b** was obtained from the AMDase IPLM-catalyzed conversion of 2-ethyl-2-phenyl malonate (**1a**). Method: GC-FID_M1. >99% ± 0.0 ee (R).*

Conversion of **1a** by AMDase VCPLLG

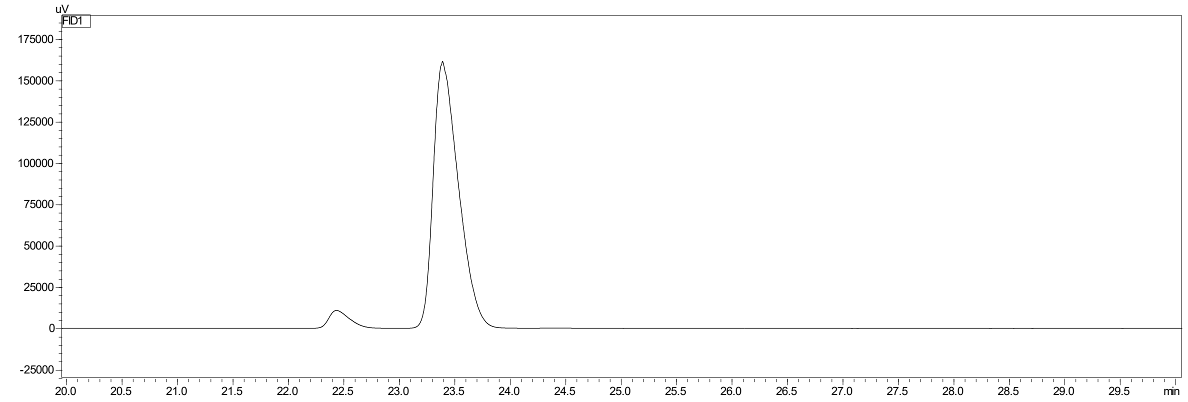

| Peak | Retention time (min) | Area |
|---|---|---|
| (*S*)-Me-**1b** | 23.4 | 2543074 |
| (*R*)-Me-**1b** | 24.2 | *not detected* |

***Figure S13.*** *Chiral GC chromatogram of methyl 2-phenyl butanoate (Me-**1b**). **1b** was obtained from the AMDase VCPLLG-catalyzed conversion of 2-ethyl-2-phenyl malonate (**1a**). Method: GC-FID_M1. >99% ± 0.0 ee (S).*

#### 5.2 2-Ethyl-3-butenoic acid (2b)

Racemic 2-ethyl-3-butenoic acid (**2b**)

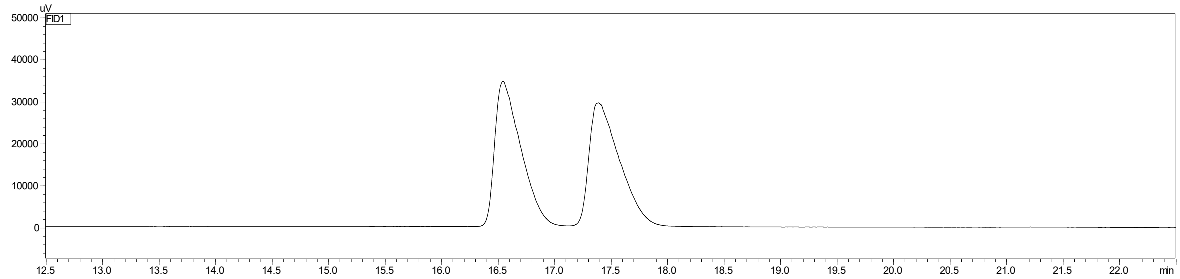

***Figure S14.*** *Chiral GC chromatogram of commercially available racemic 2-ethyl-3-butenoic acid (**2b**). Retention times of (S)-**2b** = 16.5 min and (R)-**2b** = 17.4 min. Method: GC-FID_M2.*

Conversion of **2a** by AMDase IPLL

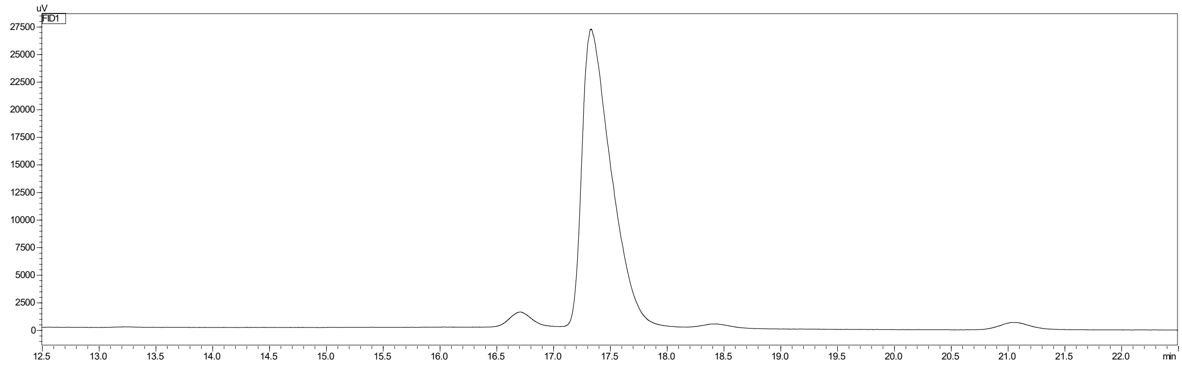

| **Peak** | **Retention time (min)** | **Area** |
|---|---|---|
| (*S*)-**2b** | 16.5 | 18640 |
| (*R*)-**2b** | 17.4 | 482104 |

***Figure S15.*** *Chiral GC chromatogram of 2-ethyl-3-butenoic acid (**2b**), obtained from the AMDase IPLL-catalyzed conversion of 2-ethyl-2-vinyl malonate (**2a**). Method: GC-FID_M2. 94.4% ± 0.8 ee (R).*

Conversion of **2a** by AMDase VPLL

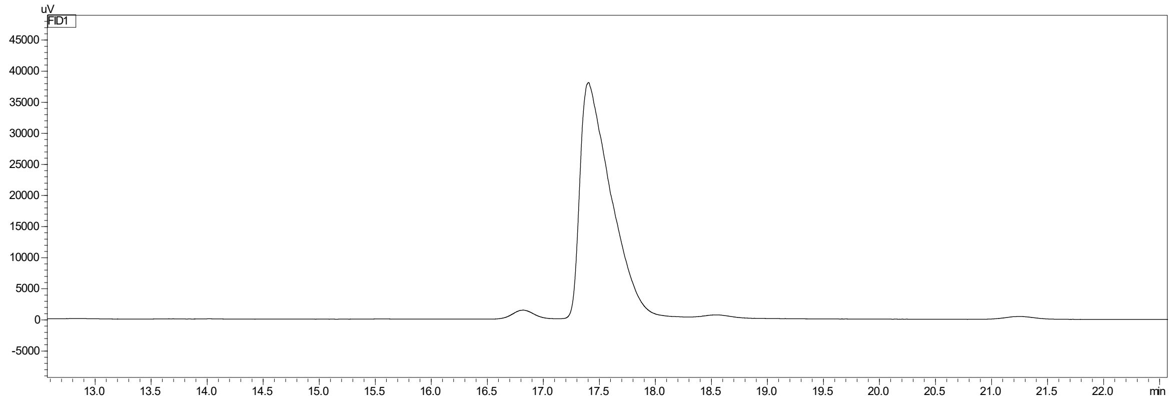

| **Peak** | **Retention time (min)** | **Area** |
|---|---|---|
| (*S*)-**2b** | 16.7 | 18384 |
| (*R*)-**2b** | 17.4 | 725262 |

***Figure S16.*** *Chiral GC chromatogram of 2-ethyl-3-butenoic acid (**2b**), obtained from the AMDase VPLL-catalyzed conversion of 2-ethyl-2-vinyl malonate (**2a**). Method: GC-FID_M2. 96.4% ± 0.6 ee (R).*

#### 5.3 Cyclohex-2-ene-1-carboxylic acid (3b)

Racemic cyclohex-2-ene-1-carboxylic acid (**3b**)

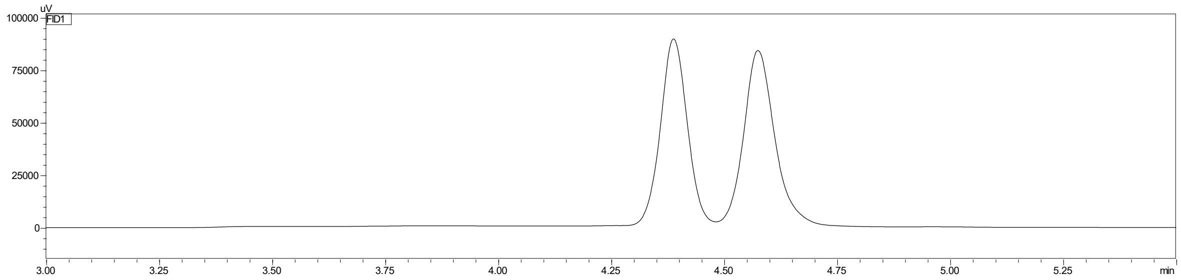

***Figure S17.*** *Chiral GC chromatogram of chemically decarboxylated racemic 2-cyclohexene-1-carboxylic acid (**3b**). Retention times of (-)-**3b** = 4.4 min and (+)-**3b** = 4.6 min. Method: GC-FID_M3.*

Conversion of **3a** by AMDase IPLL

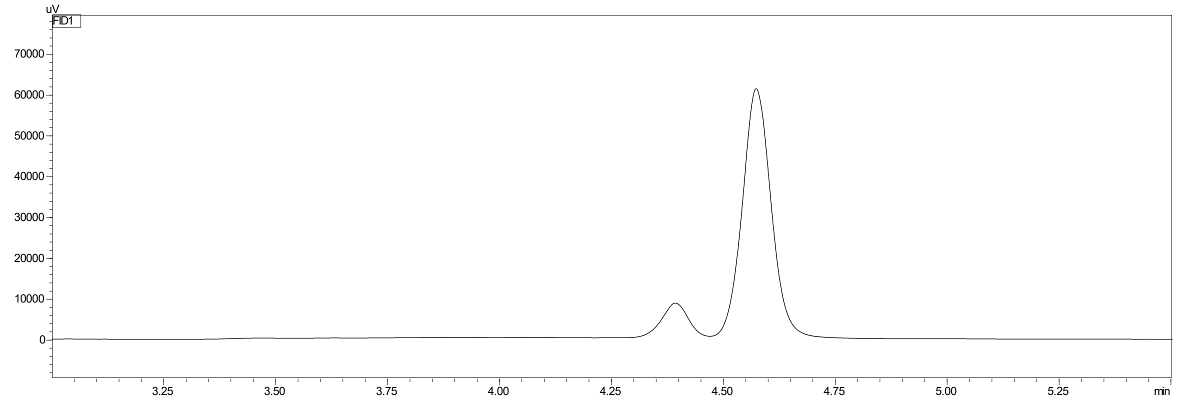

| Peak | Retention time (min) | Area |
|---|---|---|
| (-)-**3b** | 4.4 | 36018 |
| (+)-**3b** | 4.6 | 277041 |

***Figure S18.*** *Chiral GC chromatogram of 2-cyclohexene-1-carboxylic acid (**3b**), obtained from the AMDase IPLL-catalyzed conversion of 2-cyclohexene-1,1-dicarboxylic acid (**3a**). Method: GC-FID_M3. 77.0% ± 0.0 ee (+)-**3b**.*

Conversion of **3a** by AMDase VPLL

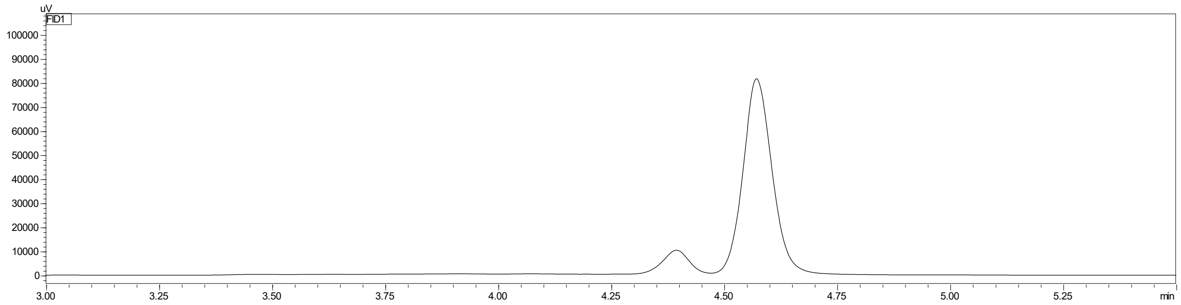

| Peak | Retention time (min) | Area |
|---|---|---|
| (-)-**3b** | 4.4 | 41663 |
| (+)-**3b** | 4.6 | 366089 |

***Figure S19.*** *Chiral GC chromatogram of 2-cyclohexene-1-carboxylic acid (**3b**), obtained from the AMDase VPLL-catalyzed conversion of 2-cyclohexene-1,1-dicarboxylic acid (**3a**). Method: GC-FID_M3. 79.0% ± 0.7 ee (+)-**3b**.*

Conversion of **3a** by AMDase VCPLLG

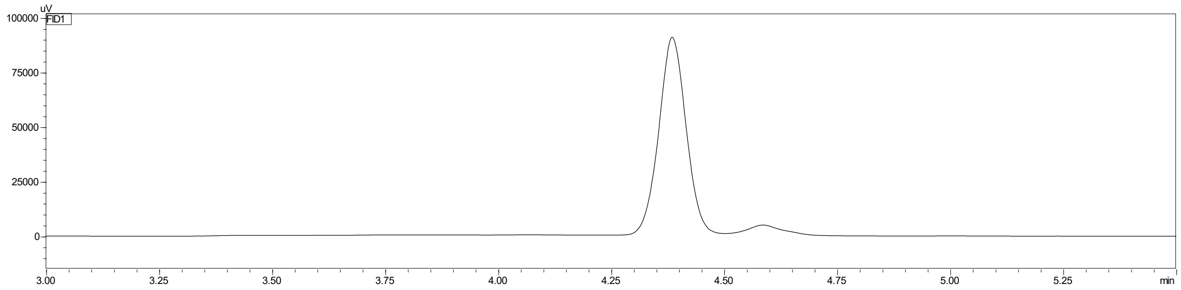

| Peak | Retention time (min) | Area |
|---|---|---|
| (-)-**3b** | 4.4 | 386298 |
| (+)-**3b** | 4.6 | 27594 |

***Figure S20.*** *Chiral GC chromatogram of 2-cyclohexene-1-carboxylic acid (**3b**), obtained from the AMDase VCPLLG-catalyzed conversion of 2-cyclohexene-1,1-dicarboxylic acid (**3a**). Method: GC-FID_M3. 86.7% ± 0.5 ee (-)-**3b**.*

#### 5.4 Methyl 2-phenyl propanoate (Me-7b)

Racemic methyl 2-phenyl propanoate (Me-**7b**)

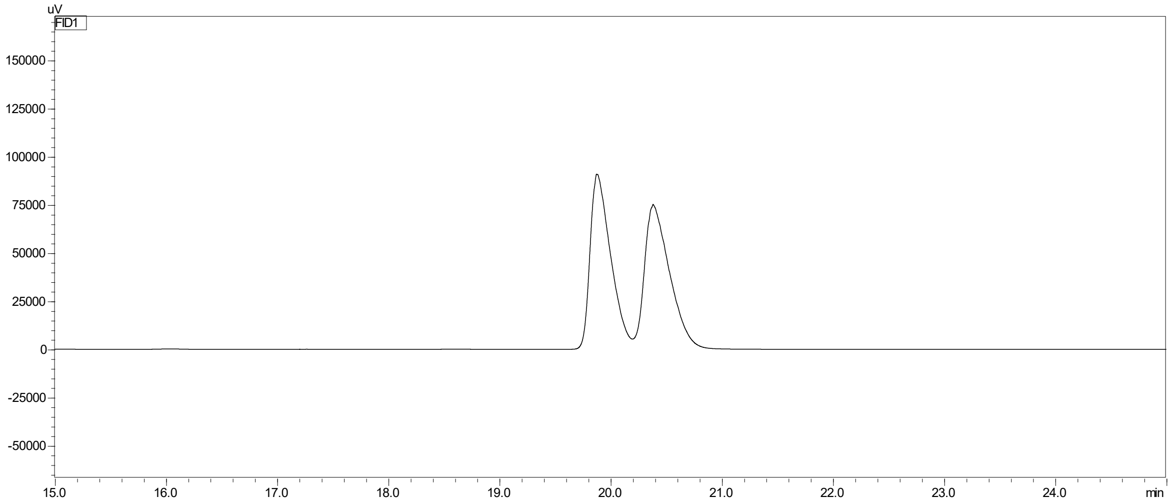

***Figure S21.*** *Chiral GC chromatogram of commercially available racemic methyl 2-phenyl propanoic acid after derivatization with $TMSCHN_2$ (Me-**7b**). Retention times of (S)-Me-**7b** = 20.0 min and (R)-Me-**7b** = 20.5 min. Method: GC-FID_M4.*

Conversion of **7a** by AMDase IPLL

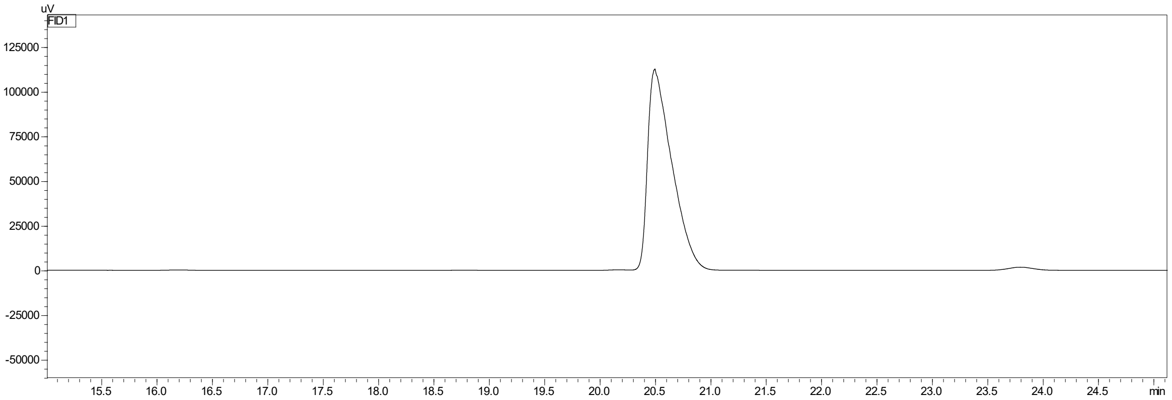

| **Peak** | **Retention time (min)** | **Area** |
|---|---|---|
| (*S*)-Me-**7b** | 20.2 | 2214 |
| (*R*)-Me-**7b** | 20.5 | 1703723 |

***Figure S22.*** *Chiral GC chromatogram of methyl 2-phenyl propanoate (Me-**7b**). **7b** was obtained from the AMDase IPLL-catalyzed conversion of 2-methyl-2-phenyl malonate (**7a**). Method: GC-FID_M4. >99% ± 0.1 ee (R).*

Conversion of **3a** by AMDase VPLL

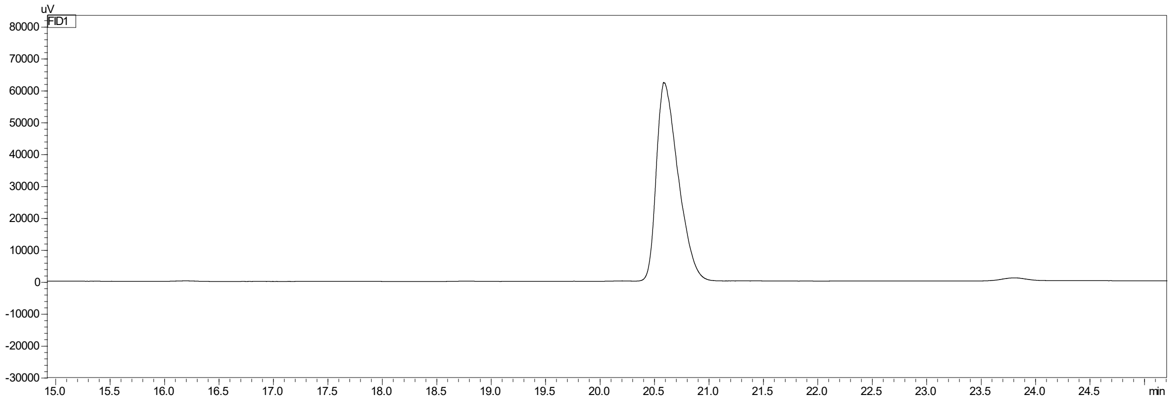

| **Peak** | **Retention time (min)** | **Area** |
|---|---|---|
| (*S*)-Me-**7b** | 20.2 | *not detected* |
| (*R*)-Me-**7b** | 20.5 | 863686 |

***Figure S23.*** *Chiral GC chromatogram of methyl 2-phenyl propanoate (Me-**7b**). **7b** was obtained from the AMDase VPLL-catalyzed conversion of 2-methyl-2-phenyl malonate (**7a**). Method: GC-FID_M4. >99% ± 0.0 ee (R).*

Conversion of **7a** by AMDase IALL

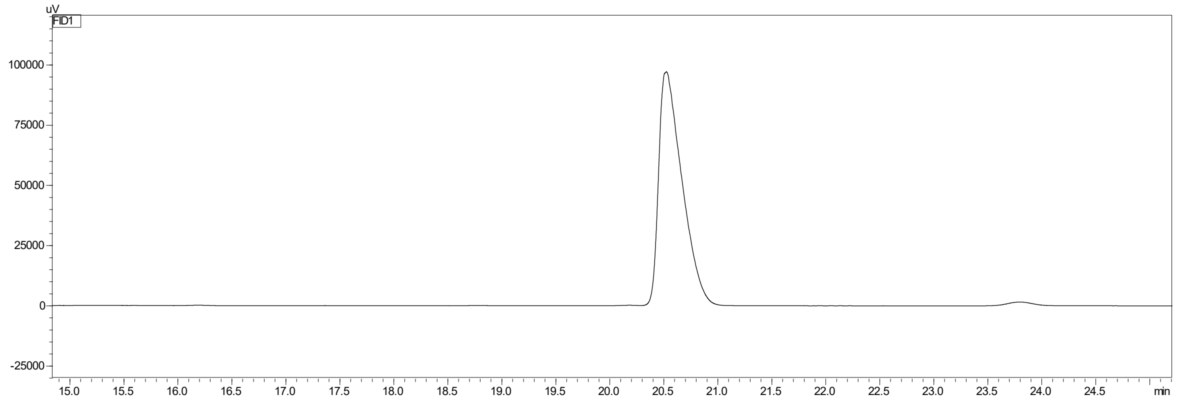

| Peak | Retention time (min) | Area |
|---|---|---|
| (*S*)-Me-**7b** | 20.2 | 1529 |
| (*R*)-Me-**7b** | 20.5 | 1457143 |

***Figure S24.*** *Chiral GC chromatogram of methyl 2-phenyl propanoate (Me-**7b**). **7b** was obtained from the AMDase IALL-catalyzed conversion of 2-methyl-2-phenyl malonate (**7a**). Method: GC-FID_M4. >99% ± 0.0 ee (R).*

Conversion of **7a** by AMDase IPVL

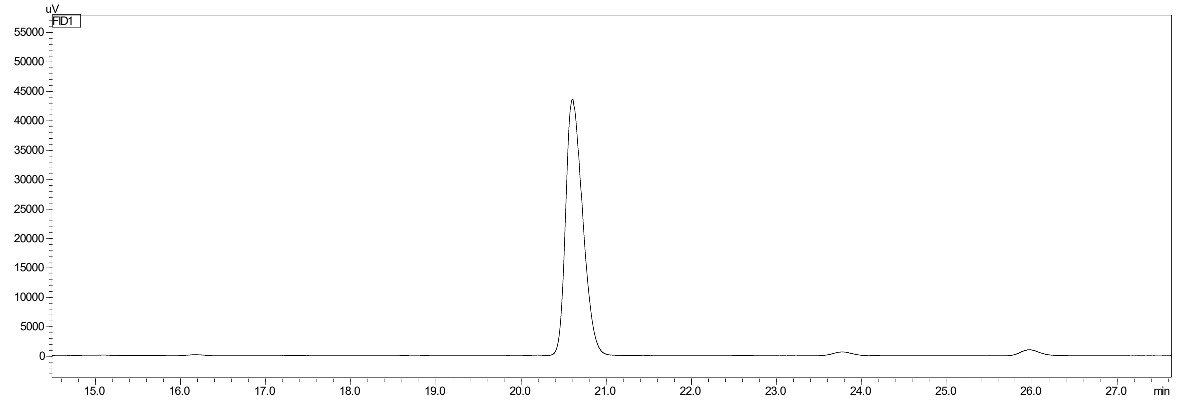

| Peak | Retention time (min) | Area |
|---|---|---|
| (*S*)-Me-**7b** | 20.2 | *not detected* |
| (*R*)-Me-**7b** | 20.5 | 603418 |

***Figure S25.*** *Chiral GC chromatogram of methyl 2-phenyl propanoate (Me-**7b**). **7b** was obtained from the AMDase IPVL-catalyzed conversion of 2-methyl-2-phenyl malonate (**7a**). Method: GC-FID_M4. >99% ± 0.0 ee (R).*

Conversion of **7a** by AMDase IPLM

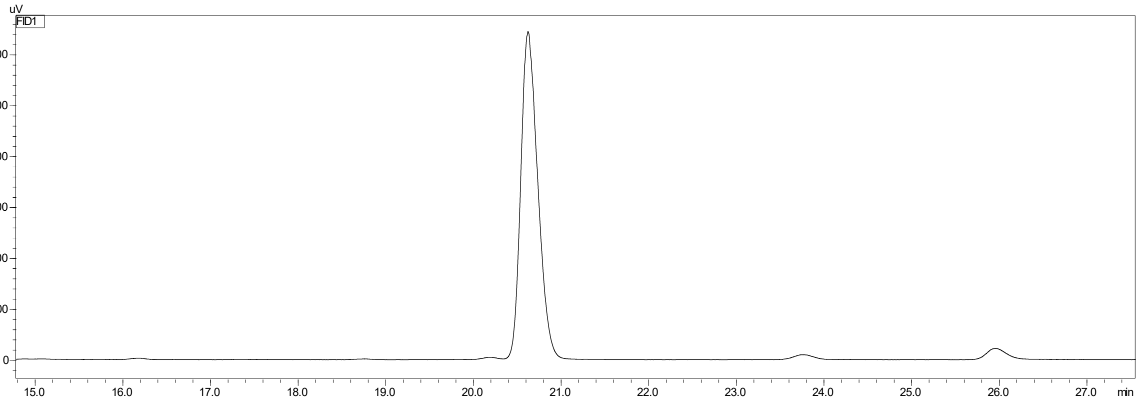

| Peak | Retention time (min) | Area |
|---|---|---|
| (*S*)-Me-**7b** | 20.2 | 197 |
| (*R*)-Me-**7b** | 20.5 | 431913 |

***Figure S26.*** *Chiral GC chromatogram of methyl 2-phenyl propanoate (Me-**7b**). **7b** was obtained from the AMDase IPLM-catalyzed conversion of 2-methyl-2-phenyl malonate (**7a**). Method: GC-FID_M4. >99% ± 0.0 ee (R).*

Conversion of **7a** by AMDase ICPLLG

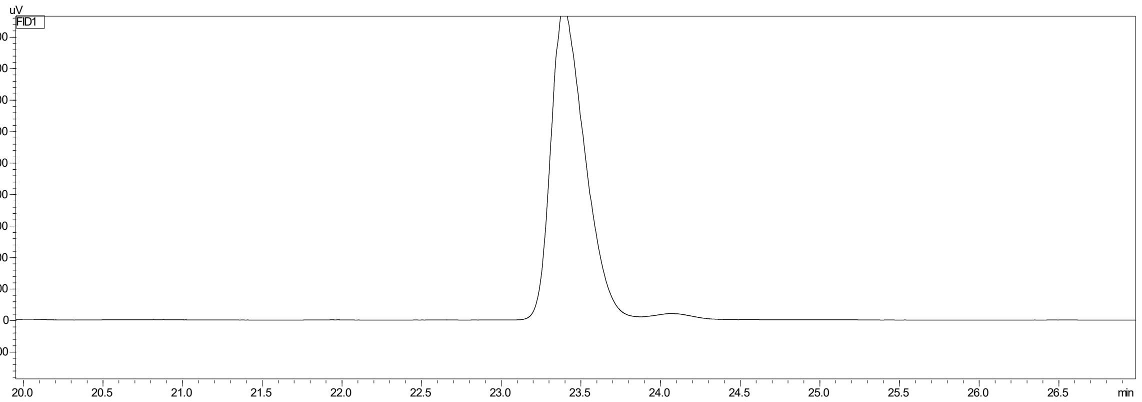

| Peak | Retention time (min) | Area |
|---|---|---|
| (*S*)-Me-**7b** | 23.4 | 735577 |
| (*R*)-Me-**7b** | 24.1 | 16395 |

***Figure S27.*** *Chiral GC chromatogram of methyl 2-phenyl propanoate (Me-**7b**). **7b** was obtained from the AMDase ICPLLG-catalyzed conversion of 2-methyl-2-phenyl malonate (**7a**). Method: GC-FID_M4. 89.0% ± 1.3 ee (S). Retention times differ due to column exchange.*

Conversion of **7a** by AMDase VCPLLG

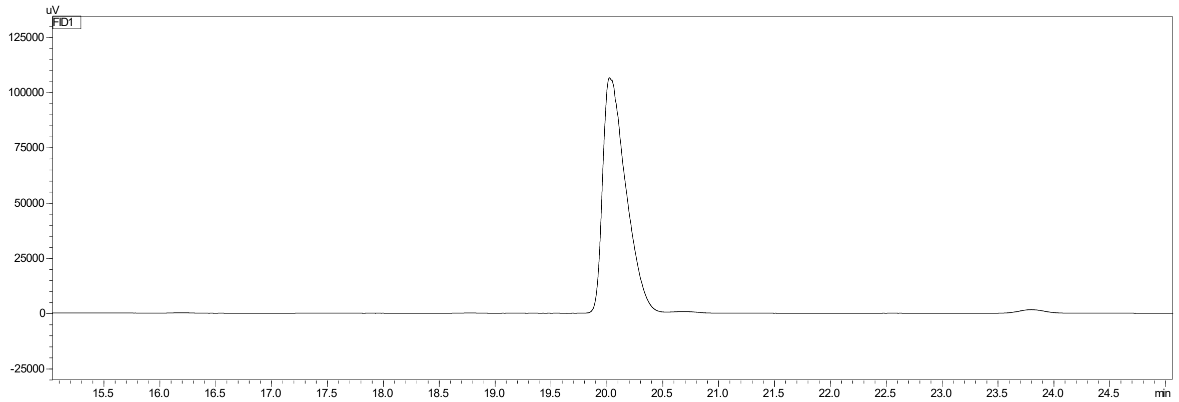

| Peak | Retention time (min) | Area |
|---|---|---|
| (*S*)-Me-**7b** | 20.2 | 1527890 |
| (*R*)-Me-**7b** | 20.7 | 4372 |

***Figure S28.*** *Chiral GC chromatogram of methyl 2-phenyl propanoate (Me-**7b**). **7b** was obtained from the AMDase VCPLLG-catalyzed conversion of 2-methyl-2-phenyl malonate (**7a**). Method: GC-FID_M4. >99.0% ± 0.1 ee (S).*

#### 5.5 2-Methyl-3-butenoic acid (8b)

Racemic 2-methyl-3-butenoic acid (**8b**)

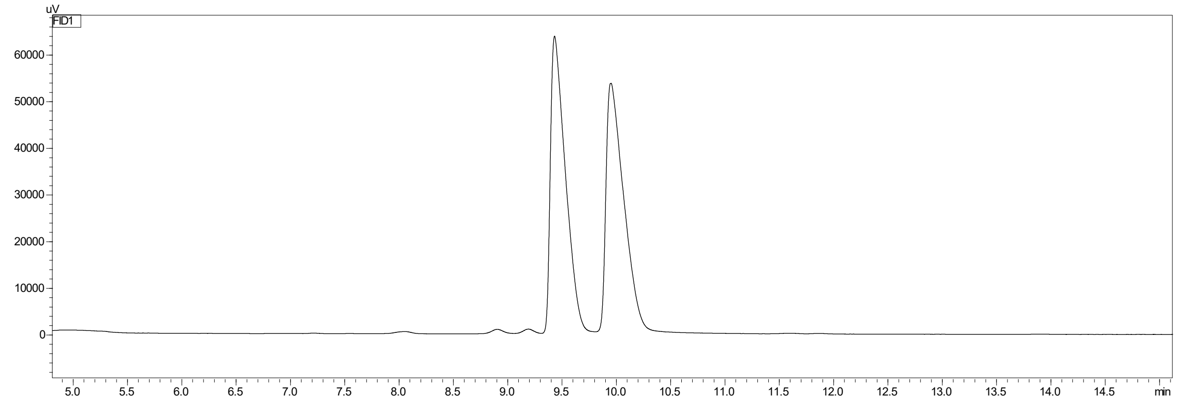

***Figure S29.*** *Chiral GC chromatogram of commercially available racemic 2-methyl-3-butenoic acid (**8b**). Retention times of (S)-**8b** = 9.5 min and (R)-**8b** = 9.9 min. Method: GC-FID_M2.*

Conversion of **8a** by AMDase VPLL

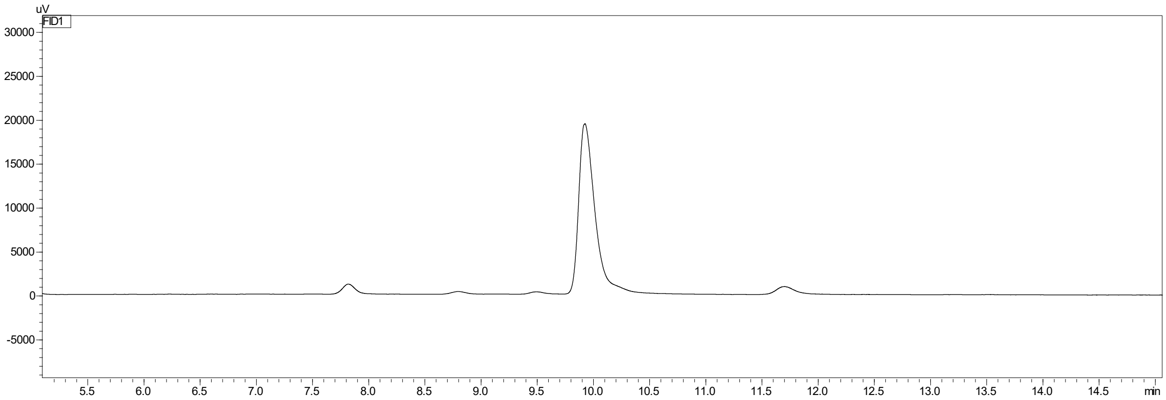

| Peak | Retention time (min) | Area |
|---|---|---|
| (*S*)-**8b** | 9.5 | 2076 |
| (*R*)-**8b** | 9.9 | 195776 |

***Figure S30.*** *Chiral GC chromatogram of 2-methyl-3-butenoic acid (**8b**), obtained from the AMDase VPLL-catalyzed conversion of 2-methyl-2-vinyl malonate (**8a**). Method: GC-FID_M2. 98% ± 0.4 ee (R).*

Conversion of **8a** by AMDase IALL

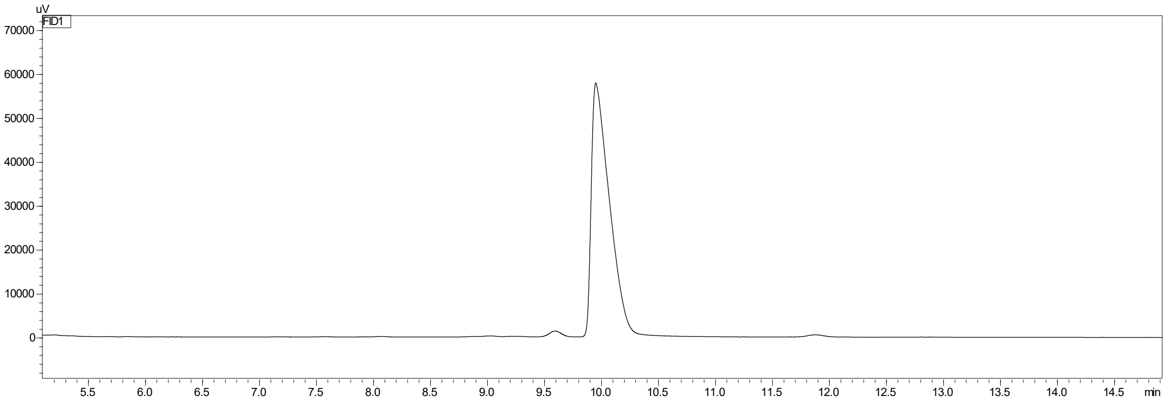

| Peak | Retention time (min) | Area |
|---|---|---|
| (*S*)-**8b** | 9.5 | 9977 |
| (*R*)-**8b** | 9.9 | 639564 |

***Figure S31.*** *Chiral GC chromatogram of 2-methyl-3-butenoic acid (**8b**), obtained from the AMDase IALL-catalyzed conversion of 2-methyl-2-vinyl malonate (**8a**). Method: GC-FID_M2. 98% ± 0.3 ee (R).*

Conversion of **8a** by AMDase IPVL

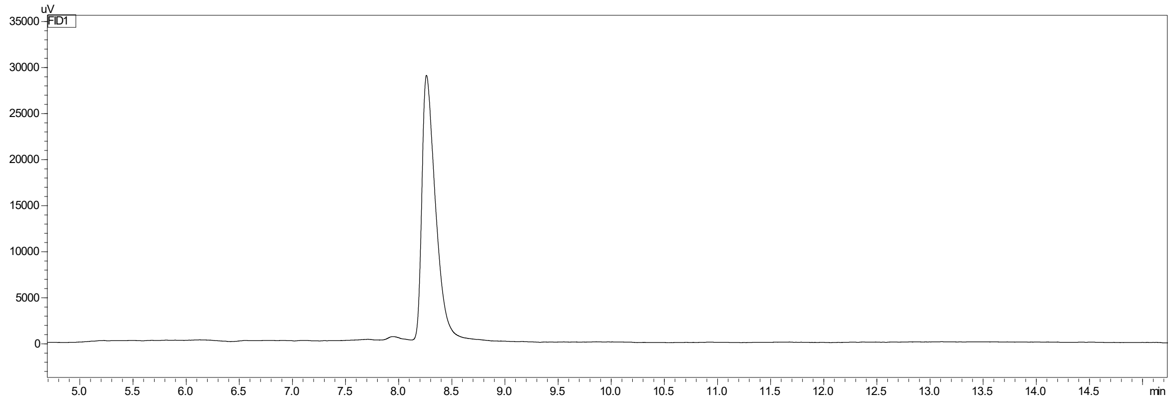

| Peak | Retention time (min) | Area |
|---|---|---|
| (*S*)-**8b** | 8.0 | 2831 |
| (*R*)-**8b** | 8.3 | 258657 |

***Figure S32.*** *Chiral GC chromatogram of 2-methyl-3-butenoic acid (**8b**), obtained from the AMDase IPVL-catalyzed conversion of 2-methyl-2-vinyl malonate (**8a**). Method: GC-FID_M2. 97.5% ± 0.3 ee (R). Retention times differ due to column exchange.*

Conversion of **8a** by AMDase IPLM

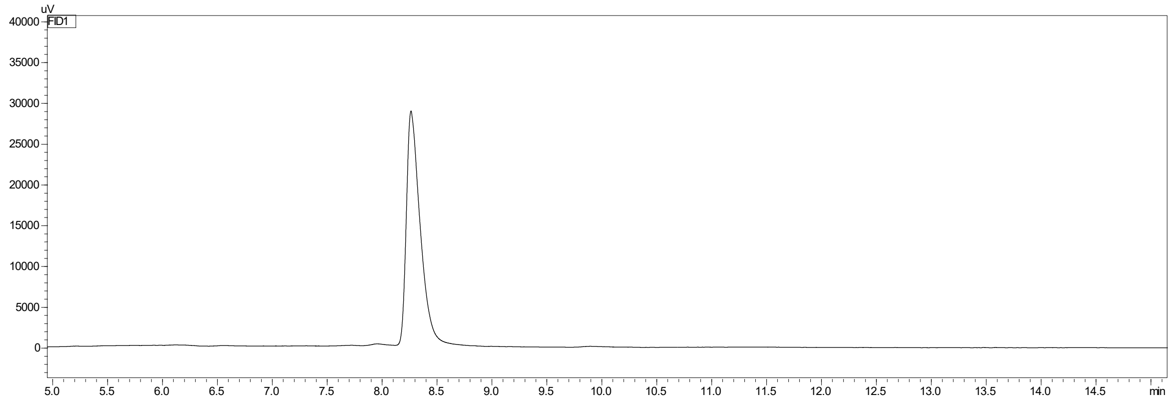

| Peak | Retention time (min) | Area |
|---|---|---|
| (*S*)-**8b** | 8.0 | 2011 |
| (*R*)-**8b** | 8.3 | 253323 |

***Figure S33.*** *Chiral GC chromatogram of 2-methyl-3-butenoic acid (**8b**), obtained from the AMDase IPLM-catalyzed conversion of 2-methyl-2-vinyl malonate (**8a**). Method: GC-FID_M2. 98.5% ± 0.0 ee (R). Retention times differ due to column exchange.*

### 6. Chiral HPLC chromatograms

#### 6.1 Ethyl-naproxen (4b)

Racemic ethyl-naproxen (**4b**)

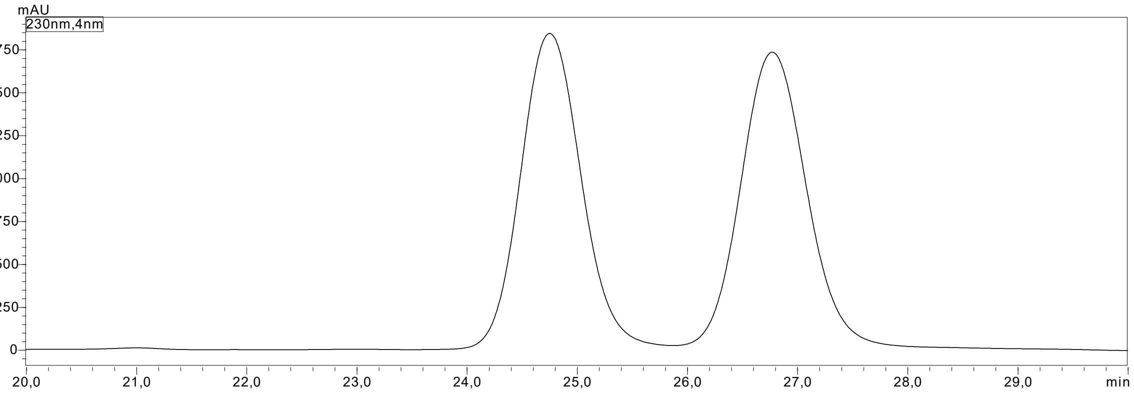

***Figure S34.*** *Chiral HPLC chromatogram of racemic ethyl-naproxen (**4b**) obtained by thermal spontaneous decarboxylation at 70°C. Retention times of (R)-**4b** = 25.1 min and (S)-**4b** = 27.2 min. Method: Chiral HPLC method.*

Conversion of **4a** by AMDase VPLL

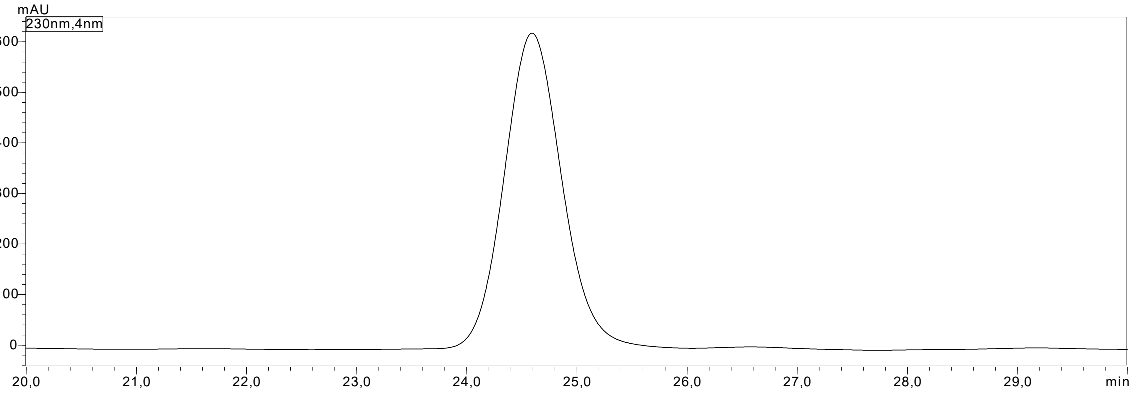

| **Peak** | **Retention time (min)** | **Area** |
|---|---|---|
| (*R*)-**4b** | 25.1 | 16162475 |
| (*S*)-**4b** | 27.1 | 27415 |

***Figure S35.*** *Chiral HPLC chromatogram of ethyl-naproxen (**4b**), obtained from the AMDase VPLL-catalyzed conversion of ethyl-naproxen malonate (**4a**). Method: Chiral HPLC method. >99% ee (R).*

Conversion of **4a** by AMDase ICPLLG

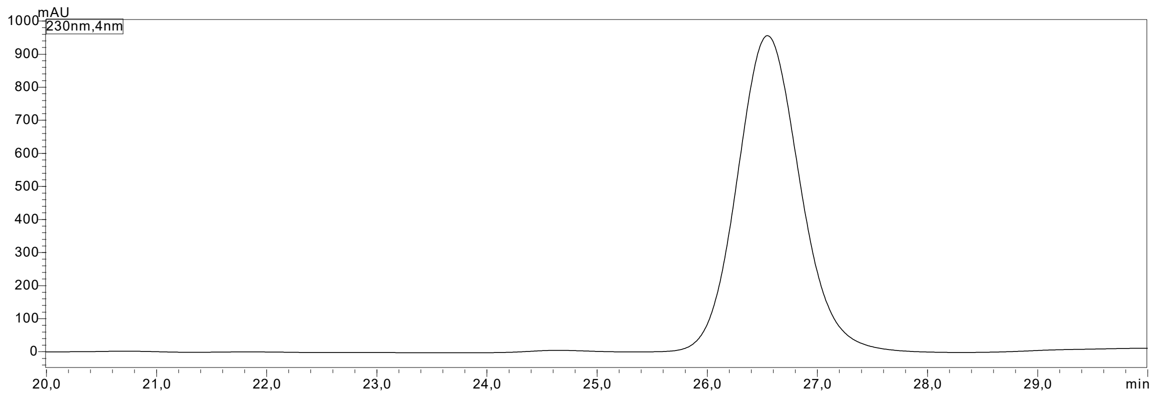

| Peak | Retention time (min) | Area |
|---|---|---|
| (*R*)-**4b** | 25.1 | 86125 |
| (*S*)-**4b** | 27.1 | 16325872 |

***Figure S36.*** *Chiral HPLC chromatogram of ethyl-naproxen (**4b**), obtained from the AMDase ICPLLG-catalyzed conversion of ethyl-naproxen malonate (**4a**). Method: Chiral HPLC method. 99% ee (S).*

#### 6.2 *n*-Propyl-naproxen (5b)

Racemic *n*-propyl-naproxen (**5b**)

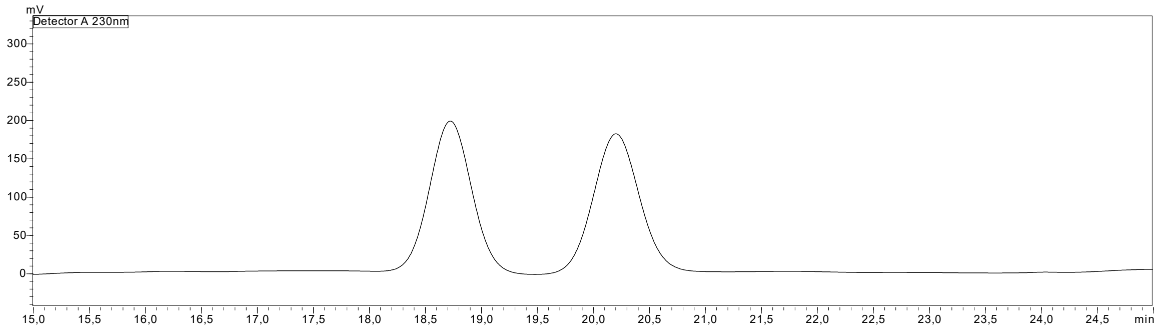

***Figure S37.*** *Chiral HPLC chromatogram of racemic n-propyl-naproxen (**5b**) obtained by thermal spontaneous decarboxylation at 70°C. Retention times: first eluting peak = 18.7min; second eluting peak = 20.2 min. Method: Chiral HPLC method. *Measurements were carried out on an HPLC PhoenixSLC-40 system without an oven, which resulted in peak shifts compared to measurements performed on the HPLC Shimadzu SLC-40 system.*

Conversion of **5a** by AMDase VPLL

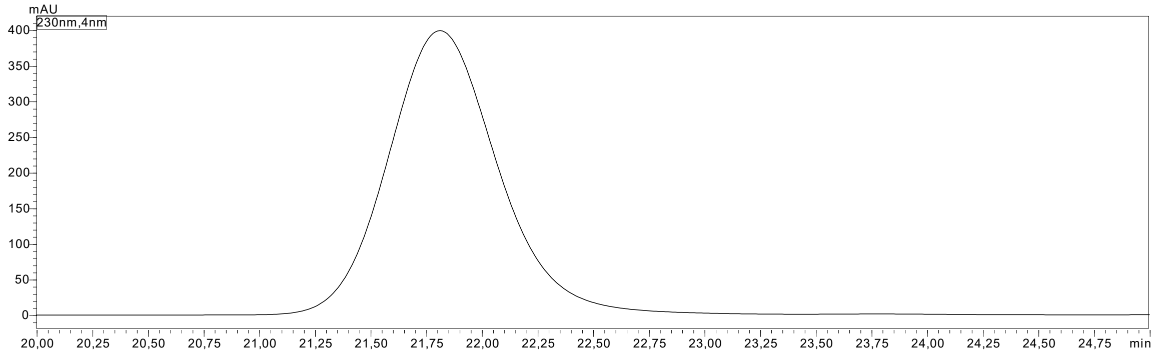

| **Peak** | **Retention time (min)** | **Area** |
|---|---|---|
| *First eluting peak* | 21.8 | 13182332 |
| *Second eluting peak* | 23.8 | 22152 |

***Figure S38.*** *Chiral HPLC chromatogram of n-propyl-naproxen (**5b**), obtained from the AMDase VPLL-catalyzed conversion of n-propyl-naproxen malonate (**5a**). Method: Chiral HPLC method. >99% ee first eluting peak.*

Conversion of **5a** by AMDase ICPLLG

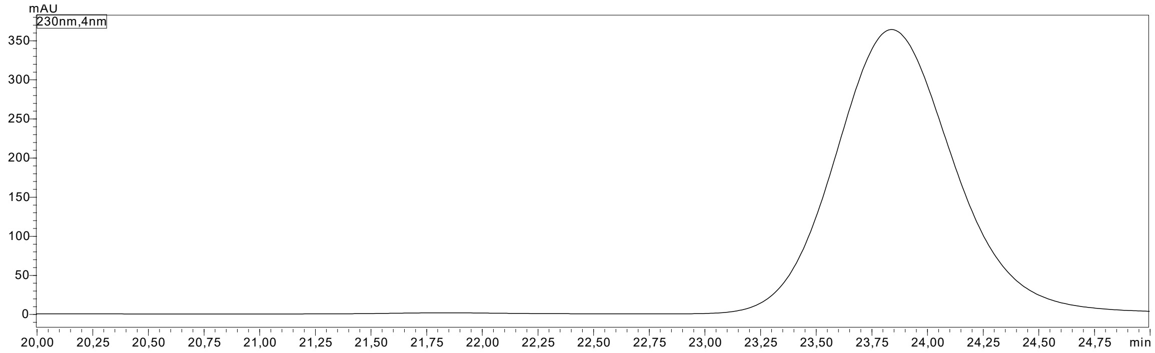

| **Peak** | **Retention time (min)** | **Area** |
|---|---|---|
| *First eluting peak* | 21.8 | 37909 |
| *Second eluting peak* | 23.8 | 15114348 |

***Figure S39.*** *Chiral HPLC chromatogram of n-propyl-naproxen (**5b**), obtained from the AMDase ICPLLG-catalyzed conversion of n-propyl-naproxen malonate (**5a**). Method: Chiral HPLC method. 99% ee second eluting peak.*

#### 7. Synthetic procedures

The substrates **1a**-**5a**, **7a**, and **8a** were synthesized according to protocols reported by van der Pol *et al.* and used for both studies.[5]

##### 7.1 Synthesis of Diethyl-2-*n*-propyl-2-vinyl-malonate (6)

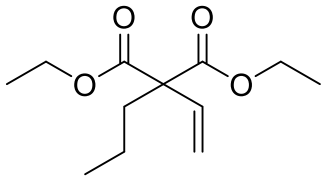

**6**

Diethyl-2-ethylidenemalonate (600 μL, 3.3 mmol) was dissolved in anhydrous DMF (30 mL) in a 50 mL Schlenk flask under inert conditions and cooled to 0 °C (ice bath). LiHMDS (4.3 mL, 1 M in THF, 4.3 mmol) was added dropwise, and the yellow solution was stirred for 30 min. After the addition of *n*-propyl iodide (0.43 mL, 4.4 mmol), the ice bath was removed, and the mixture was stirred at RT overnight. Upon complete consumption of the starting material (confirmed by GC-MS, 16 h), the reaction was quenched by the addition of sat. $NH_4Cl$ (30 mL). The product was extracted with EtOAc (3 x 30 mL). The combined organic layers were washed with 1 M LiCl (3 x 30 mL), dried over $Na_2SO_4$, filtered, and concentrated under reduced pressure. The crude product was purified *via* column chromatography (70 g $SiO_2$, cyclohexane:EtOAc = 40:1 (*v/v*)) to give the title compound as a colorless oil (380 mg, 51%).

$C_{12}H_{20}O_4$ [228.29 g·mol$^{-1}$]
$R_f$ = 0.50 (CH/EtOAc 10:1), $KMnO_4$ stain
$^1$H NMR (300 MHz, $CDCl_3$) δ 6.33 (dd, $^3J_{HH}$ = 17.8, 10.9 Hz, 1H), 5.22 (dd, $^3J_{HH}$ = 36.0, 14.3 Hz, 2H), 4.20 (q, $^3J_{HH}$ = 7.1 Hz, 4H), 2.07 – 1.96 (m, 2H), 1.25 (t, $^3J_{HH}$ = 7.1 Hz, 6H), 1.30– 1.20 (m, 2H), 0.91 (t, $^3J_{HH}$ = 7.3 Hz, 3H) ppm.

#### 7.2 Synthesis of 2-*n*-propyl-2-vinyl malonic acid (6a)

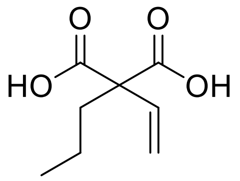

**6a**

Diethyl-2-*n*-propyl-2-vinyl-malonate (**6**, 380 mg, 1.66 mmol) was dissolved in EtOH (20 mL). The solution was cooled to 0 °C using a water-ice bath. NaOH (40%, 5 mL) was added dropwise to the solution. The reaction mixture was allowed to warm to room temperature and was stirred overnight. Upon complete consumption of the starting material (indicated by TLC, 16 h), the solvent was removed under reduced pressure. The aqueous solution was acidified with 4 M HCl to pH 4, and extracted with EtOAc (3 x 40 mL). The combined organic layers were washed with brine (40 mL), dried over $Na_2SO_4$, filtered, and concentrated under reduced pressure at 30 °C. The resulting precipitate was triturated with *n*-pentane (3 x 5 mL) and dried briefly on a rotary evaporator at 30 °C, which gave the product as an off-white powder (265 mg, 93%).

$C_8H_{12}O_4$ [172.18 g·mol$^{-1}$]

$R_f$ = 0.65 (cyclohexane:EtOAc = 4:2 (v/v); 10 drops AcOH to 6 mL eluent, $KMnO_4$)

m.p. = 108-110 °C

$^1$H NMR (300 MHz, MeOD) δ 6.33 (dd, $^3J_{HH}$ = 17.8, $^3J_{HH}$ = 10.9 Hz, 1H), 5.22 (dd, $^3J_{HH}$ = 17.8, $^3J_{HH}$ = 10.9 Hz, 2H), 2.04 – 1.91 (m, 1H), 1.37 – 1.13 (m, 2H), 0.93 (t, $^3J_{HH}$ = 7.4 Hz, 3H) ppm.

$^{13}$C NMR (76 MHz, MeOD) δ 174.0, 137.3, 116.2, 61.3, 38.2, 18.8, 14.7 ppm.

#### 9. NMR spectra

$^1$H and $^{13}$C-NMR spectra of 2-cyclohexene-carboxylic acid (**3b**), in $CDCl_3$.

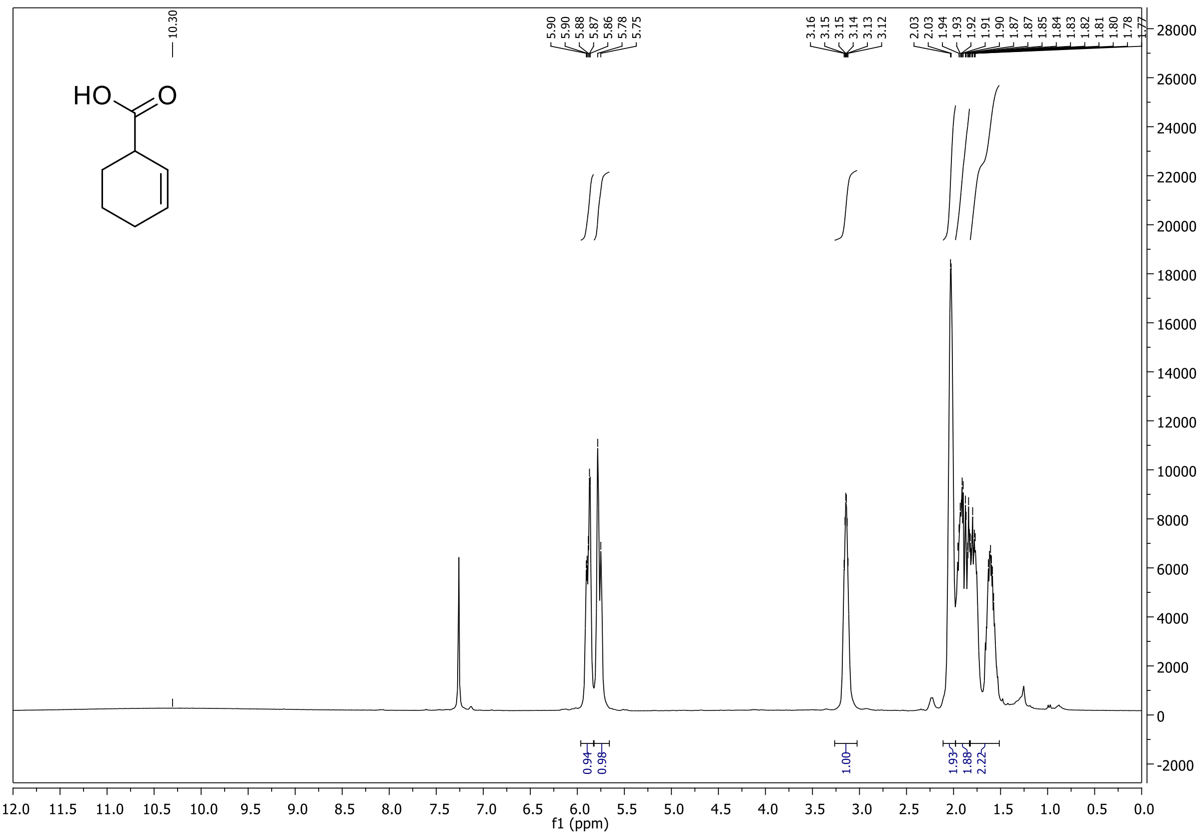

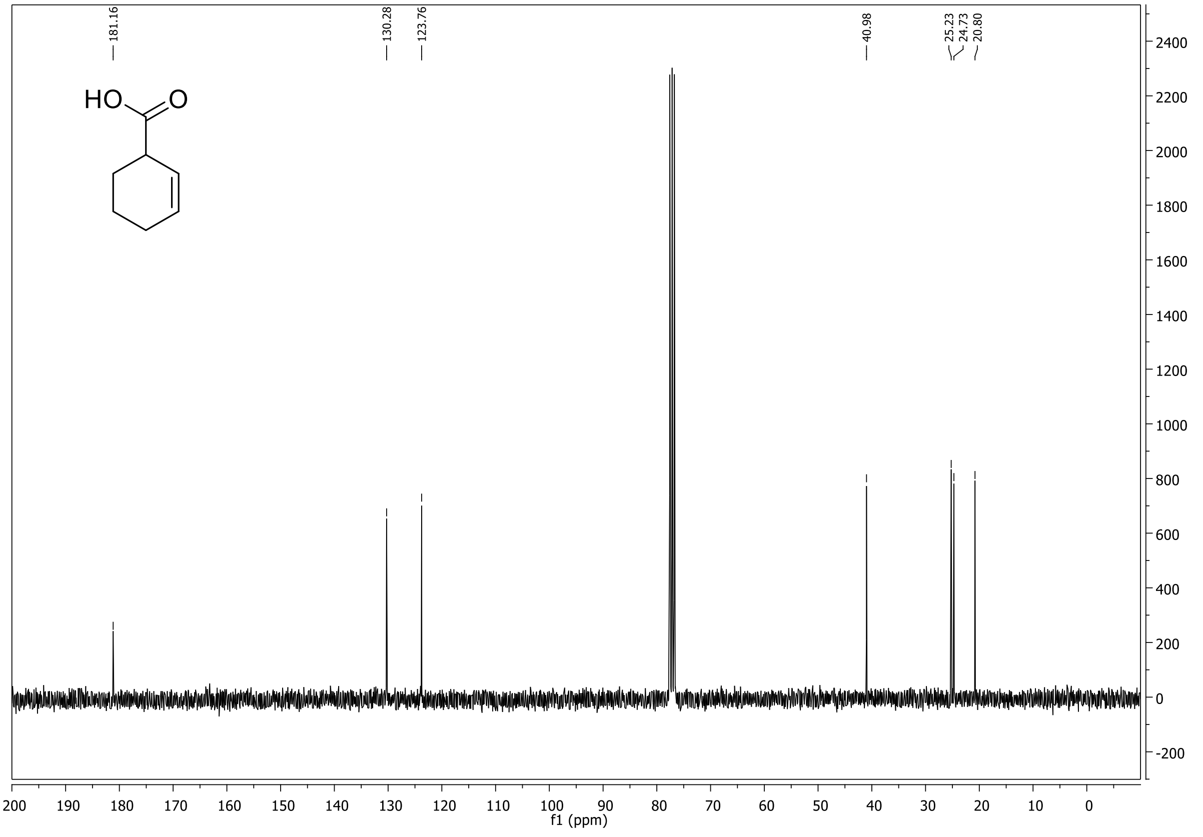

$^1$H and $^{13}$C-NMR spectra of ethyl-naproxen (**4b**), in $CDCl_3$.

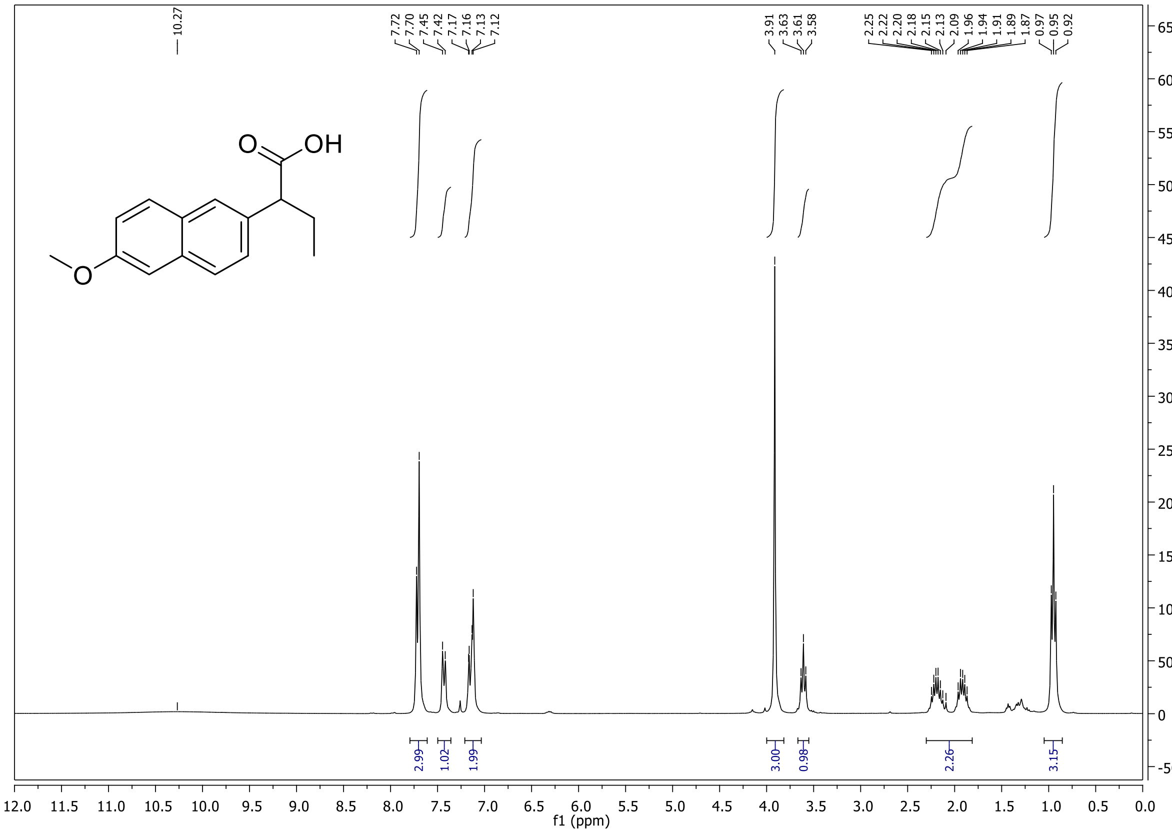

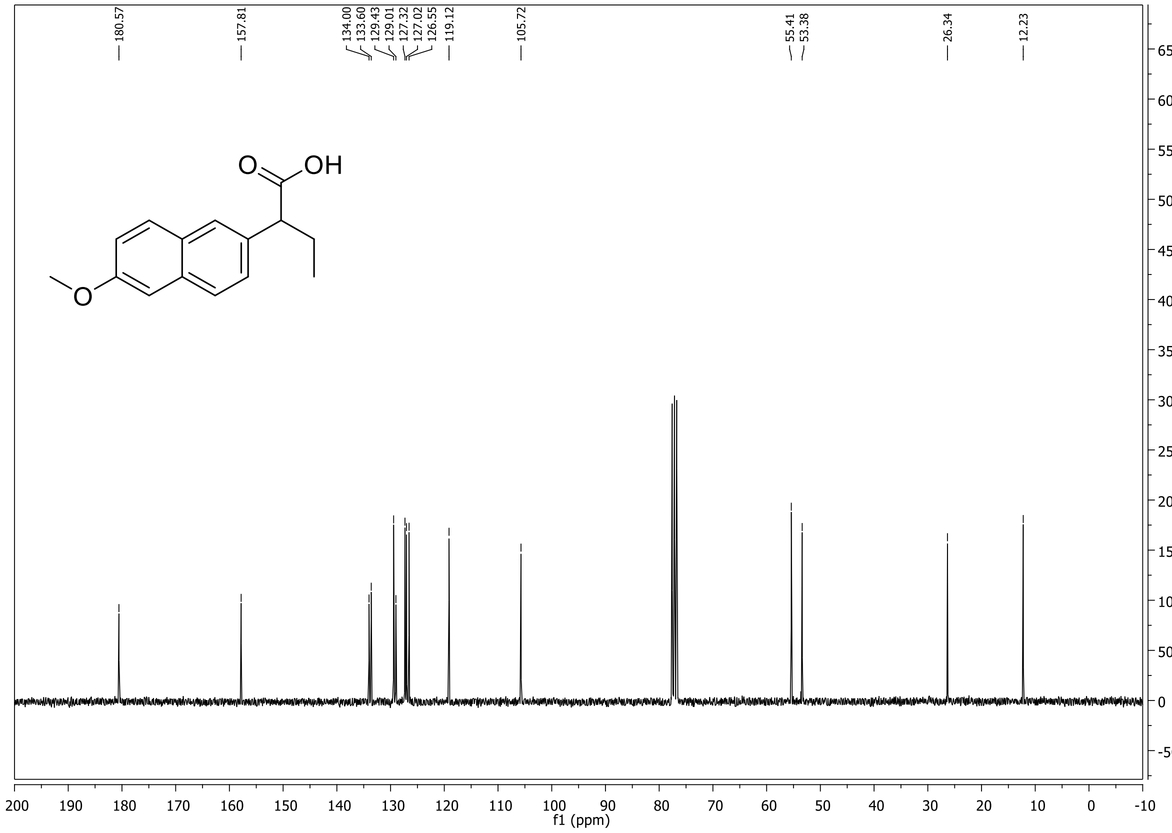

$^{1}H$-NMR spectrum of diethyl-2-*n*-propyl-2-vinyl-malonate (**6**), in $CDCl_3$.

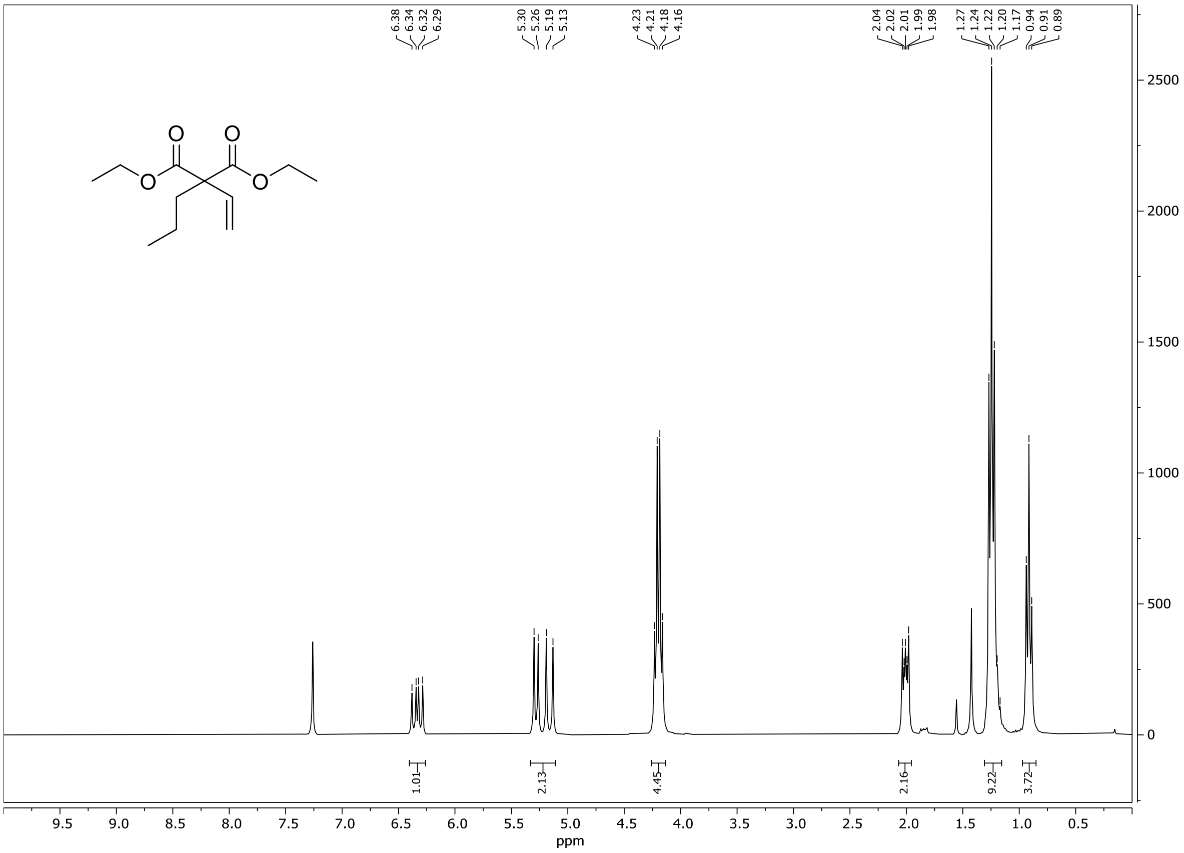

$^{1}H$ and $^{13}C$-NMR spectrum of 2-*n*-propyl-2-vinyl-malonate (**6a**), in MeOD.

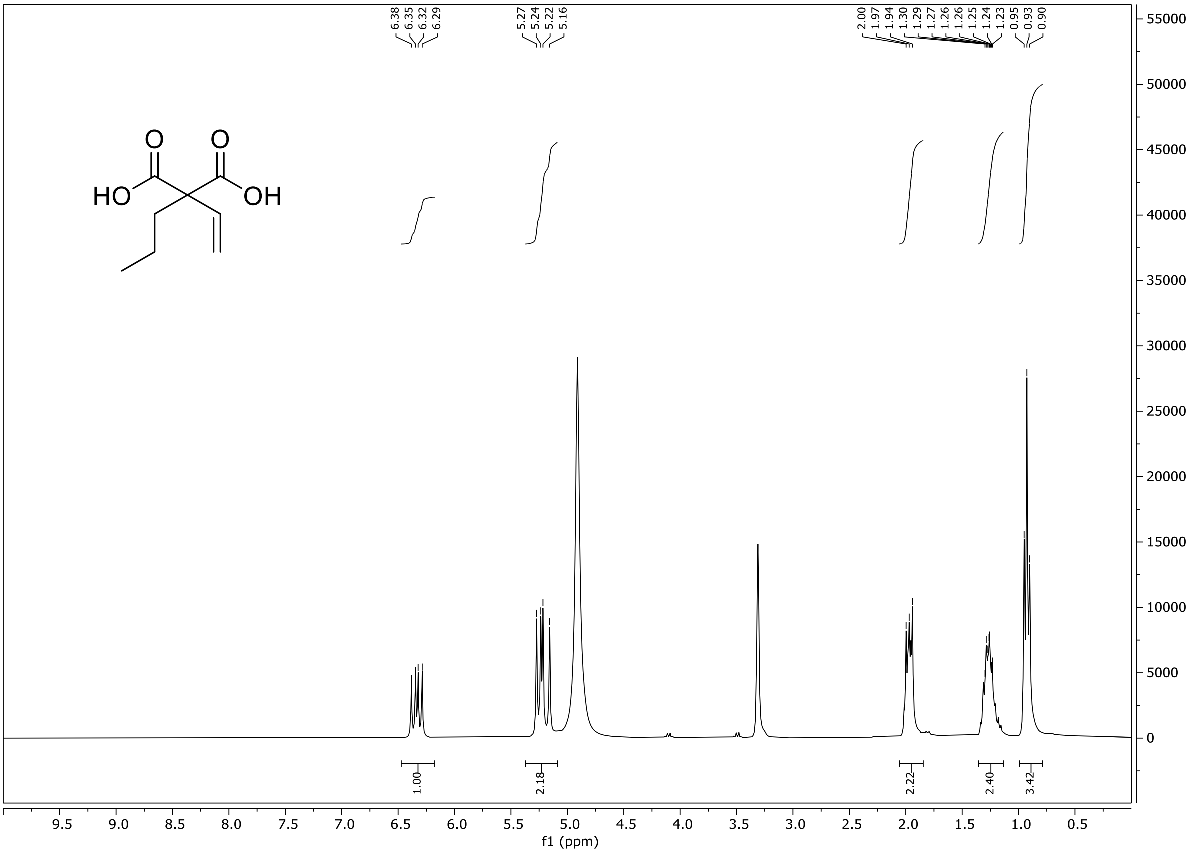

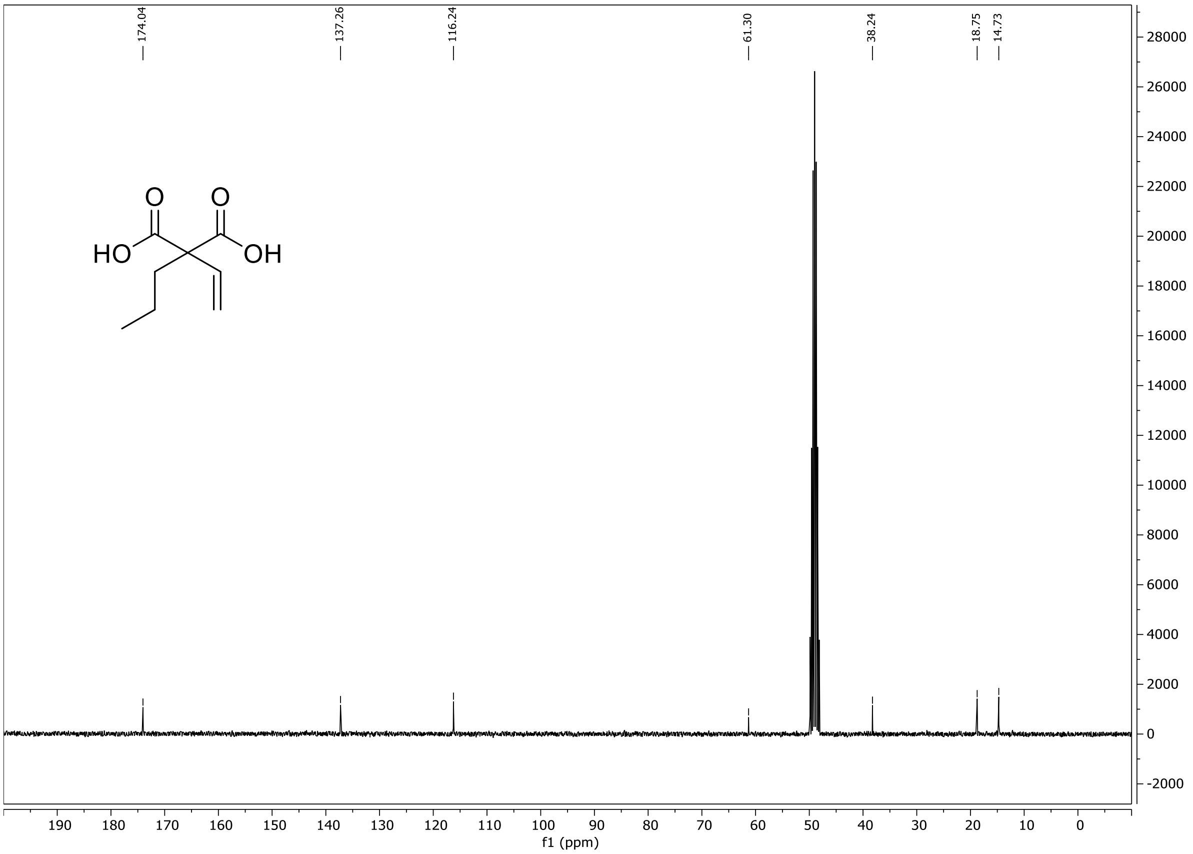
174.04
137.26
116.24
61.30
38.24
18.75
14.73
HO
OH
O
O
f1 (ppm)
190 180 170 160 150 140 130 120 110 100 90 80 70 60 50 40 30 20 10 0
28000 26000 24000 22000 20000 18000 16000 14000 12000 10000 8000 6000 4000 2000 0 -2000
